# Phytoplasma SAP11 effector destabilization of TCP transcription factors differentially impact development and defence of Arabidopsis versus maize

**DOI:** 10.1101/574319

**Authors:** Pascal Pecher, Gabriele Moro, Maria Cristina Canale, Sylvain Capdevielle, Archana Singh, Allyson MacLean, Akiko Sugio, Chih-Horng Kuo, Joao R. S. Lopes, Saskia A. Hogenhout

**Affiliations:** John Innes Centre, Department of Crop Genetics, Norwich Research Park, Norwich NR4 7UH, UK; Luiz de Queiroz College of Agriculture, Department of Entomology and Acarology, University of São Paulo, Piracicaba 13418-900, Brazil; Academia Sinica, Institute of Plant and Microbial Biology, Taipei, Taiwan; Agricultural Research Company of Santa Catarina State (Epagri), Chapeco 89809-450, Brazil; Earlham Institute, Norwich Research Park, Norwich NR4 7UZ, UK; Department of Biology, University of Ottawa, Ottawa, ON, K1N 6N5, Canada; INRA, UMR Institut de Génétique, Environnement et Protection des Plantes (IGEPP) Domaine de la Motte, 35653 Le Rheu cedex - France

**Keywords:** Insect vector, Phytoplasma, Plant architecture, Plant defence response, Plant development, Plant-insect interactions, SAP11 effectors, TCP transcription factors.

## Abstract

Phytoplasmas are insect-transmitted bacterial pathogens that colonize a wide range of plant species, including vegetable and cereal crops, and herbaceous and woody ornamentals. Phytoplasma-infected plants often show dramatic symptoms, including proliferation of shoots (witch’s brooms), changes in leaf shapes and production of green sterile flowers (phyllody). Aster Yellows phytoplasma Witches’ Broom (AY-WB) infects dicots and its effector, secreted AYWB protein 11 (SAP11), was shown to be responsible for the induction of shoot proliferation and leaf shape changes of plants. SAP11 acts by destabilizing TEOSINTE BRANCHED 1-CYCLOIDEA-PROLIFERATING CELL FACTOR (TCP) transcription factors, particularly the class II TCPs of the CYCLOIDEA/TEOSINTE BRANCHED 1 (CYC/TB1) and CINCINNATA (CIN)-TCP clades. SAP11 homologs are also present in phytoplasmas that cause economic yield losses in monocot crops, such as maize, wheat and coconut. Here we show that a SAP11 homolog of Maize Bushy Stunt Phytoplasma (MBSP), which has a range primarily restricted to maize, destabilizes only TB1/CYC TCPs. SAP11_MBSP_ and SAP11_AYWB_ both induce axillary branching and SAP11_AYWB_ also alters leaf development of *Arabidopsis thaliana* and maize. However, only in maize, SAP11_MBSP_ prevents female inflorescence development, phenocopying maize *tb1* lines, whereas SAP11_AYWB_ prevents male inflorescence development and induces feminization of tassels. SAP11_AYWB_ promotes fecundity of the AY-WB leafhopper vector on *A. thaliana* and modulates the expression of *A. thaliana* leaf defence response genes that are induced by this leafhopper, in contrast to SAP11_MBSP_. Neither of the SAP11 effectors promote fecundity of AY-WB and MBSP leafhopper vectors on maize. These data provide evidence that class II TCPs have overlapping but also distinct roles in regulating development and defence in a dicot and a monocot plant species that is likely to shape SAP11 effector evolution depending on the phytoplasma host range.

**Author summary:** Phytoplasmas are parasites of a wide range of plant species and are transmitted by sap-feeding insects, such as leafhoppers. Phytoplasma-infected plants are often easily recognized because of their dramatic symptoms, including shoot proliferations (witch’s brooms) and altered leaf shapes, leading to severe economic losses of crops, ornamentals and trees worldwide. We previously found that the virulence protein SAP11 of aster yellows witches’ broom phytoplasma (AY-WB) interferes with a specific group of plant transcription factors, named TCPs, leading to witches’ brooms and leaf shape changes of the model plant *Arabidopsis thaliana*. SAP11 has been characterized in a number of other phytoplasmas. However, it is not known how phytoplasmas and their SAP11 proteins modulate processes in crops, including cereals such as maize. We identified a SAP11 homolog in Maize bushy stunt phytoplasma (MBSP), a pathogen that can cause severe yield losses of maize. We found that SAP11 interactions with TCPs are conserved between maize and Arabidopsis, and that MBSP SAP11 interferes with less TCPs compared to AY-WB SAP11. This work provides new insights into how phytoplasmas change maize architecture and corn production. Moreover, we found that TCPs regulate leaf defence responses to phytoplasma leafhopper vectors in Arabidopsis, but not in maize.

## Introduction

Phytoplasmas (“*Candidatus* (*Ca.*) Phytoplasma”) are economically important plant pathogens that infect a broad range of plant species. The more than 1000 phytoplasmas described so far comprise three distinct clades within a monophyletic group of the class Mollicutes that are characterized by the lack of a bacterial cell wall and small genomes (580 kb to 2200 kb) [1–3]. These fastidious pathogens are restricted to the phloem sieve cells of the plant vasculature and depend on phloem-sap-feeding insect vectors, including leafhoppers, planthoppers and psyllids, for transmission and spread in nature [4]. Many phytoplasmas induce dramatic changes in plant architecture such as increased axillary branching (often referred to as witches’ broom), formation of leaf-like flowers (phyllody), the production of green floral organs such as petals and stamens (virescence), changes of leaf shape, and premature bolting [5–10].

Phytoplasmas change plant architecture via the secretion of proteinaceous effectors that interact with and destabilize plant transcription factors with fundamental roles in regulating plant development. Effectors of Aster yellows phytoplasma strain Witches Broom (AY-WB; “*Ca*. Phytoplasma asteris”) are particularly well characterized. AY-WB and its predominant leafhopper vector *Macrosteles quadrilineatus* have broad host ranges that mostly include dicots, including *Arabidopsis thaliana* [6]. SAP11 destabilizes Arabidopsis TEOSINTE BRANCHED1-CYCLOIDEA-PROLIFERATING CELL FACTOR (TCP) transcription factors, and specifically class II TCPs, leading to the induction of axillary branching and changes in leaf shape of this plant [8,11], and SAP54 degrades Arabidopsis MADS-box transcription factors leading to changes in flower development that resemble phyllody and virescence symptoms [9,12]. Moreover, both effectors modulate plant defence responses leading to increased colonization of *M. quadrilineatus* on *A. thaliana* [8,9,13]. For SAP11_AYWB_ this involves the inhibition of jasmonate (JA) synthesis [8]. SAP11 and SAP54 homologs of other phytoplasmas also target TCPs and MADS, respectively, leading to corresponding changes in plant development and architecture [10, 14–16]. The majority of phytoplasma effector genes lie within composite-transposon-like pathogenicity islands named potential mobile units (PMUs) that are prone to recombination and horizontal gene transfer [17–20].

Maize bushy stunt phytoplasma (MBSP) belongs to the Aster yellows (AY) group (16SrI) “*Ca*. P. asteris” [21] and is the only known member of this group to be largely restricted to maize (*Z. mays* L.), whereas the majority, including AY-WB, are transmitted by polyphagous insects and infect dicotyledonous plants [13,22]. MBSP is transmitted by the maize-specialist insects *Dalbulus maidis* and *D. elimatus*; both MBSP and insect vectors are thought to have co-evolved with maize since its domestication from teosinte [23]. Symptoms of MBSP-infected maize plants include the formation of long lateral branches, decline in ear development and emergence of leaves that are often twisted with ripped edges and that display chlorosis and reddening [13]. We previously identified a SAP11 homolog in the MBSP genome [22] and SAP11_MBSP_ is identical in sequence among multiple MBSP isolates collected from Mexico and Brazil [13]. *SAP11_AYWB_* and *SAP11_MBSP_* lie on microsyntenic regions within the phytoplasma genomes, indicating that these effectors are likely to have common ancestry [13]. However, *D. maidis* does not produce more progeny on MBSP-infected plants that show advanced disease symptoms; the insects prefer infected plants that are non-symptomatic [24]. In this study we wished to compare the roles of SAP11_AYWB_ and SAP11_MBSP_ in symptom induction and plant defence to insect vectors of *A. thaliana* and maize.

TCP transcription factors comprise an ancient plant-specific family [25] that are distinguished from other transcription factors by a conserved ± 60 amino acid TCP domain [26]. The TCP domain consists of a helix-loop-helix region that form TCP homo or heterodimers and a basic region that mediates interactions of TCP dimers with DNA motifs [27] and is required for SAP11 binding to TCPs [11]. TCP transcription factors are grouped into three clades based on TCP domain sequences: (i) class I PROLIFERATING CELL FACTOR-type TCPs (PCF clade); (ii) class II CINCINNATA-type TCPs (CIN clade); and (iii) class II CYCLOIDEA/TEOSINTE BRANCHED 1-type TCPs (CYC/TB1-clade) [28]. The latter is also known as the glutamic acid-cysteine-glutamic acid (ECE) clade [29]. PCFs promote cell proliferation, whereas CIN clade TCPs promote leaf and petal cell maturation and differentiation and have antagonistic roles to PCFs [30–33]. The ECE clade includes maize TEOSINTE BRANCHED 1 (TB1) and TB1 homologs of *A. thaliana* BRANCHED 1 (BRC1) and BRC2, that repress the development of axillary branches in plants [34–37], and CYCLOIDEA (CYC) that control flower symmetry [38]. TB1 and genes in the TB1 network have been targeted for selection during maize domestication from a teosinte ancestor [39,40].

Here we show that SAP11_AYWB_ and SAP11_MBSP_ have overlapping but distinct specificities for destabilizing class II TCP transcription factors. The SAP11 effectors induce unique phenotypes in Arabidopsis and maize that indicate divergent roles of class II TCP transcription factors in regulating development and defence in the two plant species. We argue that SAP11_MBSP_ evolution may be constrained due to the specific functionalities of class II TCPs in maize.

## Results

### Phytoplasma SAP11_AYWB_ binds and destabilizes both Arabidopsis CIN and CYC/TB1 TCPs and SAP11_MBSP_ only CYC/TB1 TCPs

SAP11_AYWB_ and SAP11_MBSP_ interaction specificities for Arabidopsis TCPs (AtTCPs) were investigated via yeast two-hybrid (Y2H) assays and protein destabilization assays in *A. thaliana* mesophyll protoplasts. In the protoplast experiments, SAP11_AYWB_ destabilized the majority of AtCIN-TCPs and none of the class I AtTCPs (Fig. 1A), confirming previous results [8]. In addition, SAP11_AYWB_ also destabilized CYC/TB1-TCPs BRC1 and BRC2 but not the five Arabidopsis class I TCPs (Fig. 1A). In contrast, SAP11_MBSP_ destabilized the CYC/TB1 TCPs BRC1 and BRC2, whereas 7 out of 8 class II AtCIN-TCPs and all tested class I AtTCPs remained stable (Fig. 1A). The Y2H assays showed that SAP11_AYWB_ interacts with Arabidopsis CIN-TCPs (Fig. 1B), confirming previous data [8,11], whereas SAP11_MBSP_ did not (Fig. 1B). However, both SAP11_AYWB_ and SAP11_MBSP_ interacted with CYC/TB1 BRC1 and BRC2 (Fig. 1B). Therefore, SAP11_MBSP_ binds and destabilizes a narrower set of class II TCPs compared to SAP11_AYWB_.

**Fig. 1.**
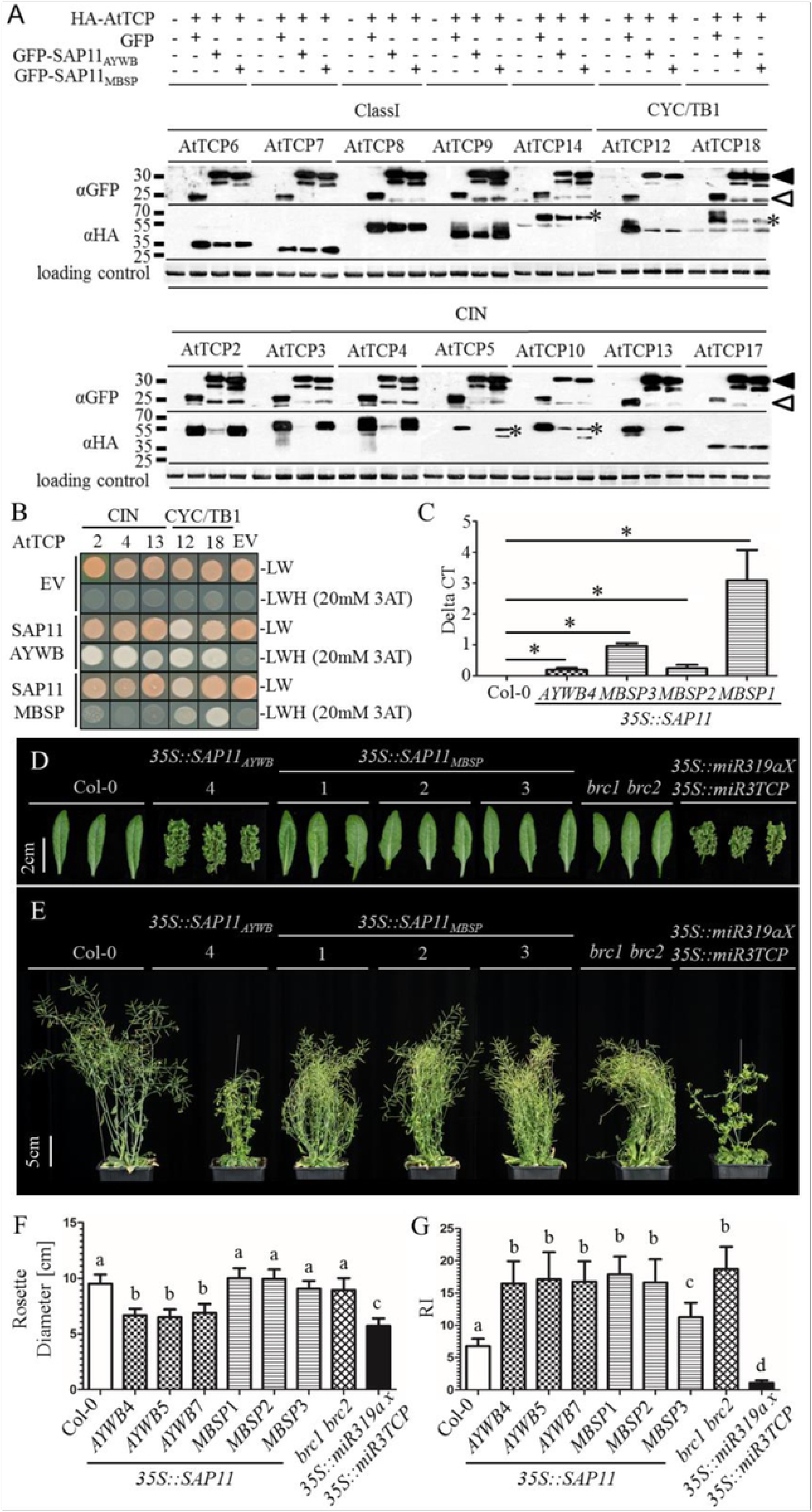
SAP11_AYWB_ and SAP11_MBSP_ interactions with *A. thaliana* TCP transcription factors. (A) Western blots of *A. thaliana* protoplast destabilization assays; SAP11_AYWB_ and SAP11_MBSP_ destabilize the CYC/TB1 TCPs BRC1 (AtTCP18) and BRC2 (AtTCP12) and SAP11_AYWB_ also all class II CIN-TCPs, whereas the SAP11 homologs did not destabilize class I TCPs. GFP-tagged SAP11 (filled arrowheads) or GFP alone (open arrowheads) and HA-tagged TCPs were detected with specific antibodies to GFP and HA, respectively, as indicated at left of the blots. *band of the correct size in case of multiple bands on the blots. Loading controls: Amidoblack-stained large RUBISCO subunit. (B) Yeast two-hybrid assays of interactions of SAP11_AYWB_ with CIN and CYC/TB1-TCPs and SAP11_MBSP_ with CYC/TB1-TCPs. Positive interactions are visible by yeast growth on SD-LWH selection media containing 20 mM 3-Amino-1,2,4-triazole (3AT). EV=empty vector controls showing absence of auto activations. (C) qRT-PCRs of transcripts of *SAP11_AYWB_* and *SAP11_MBSP_* transgenes in *A. thaliana* lines shown in D-F. *p<0.01, students t-test compared to Col-0, n=3. (D-G) *35S::SAP11_AYWB_* stable transgenic *A. thaliana* (Col-0) lines phenocopy both the *A. thaliana brc1-2 brc2-1* (*brc1 brc2*) double (Col-0) mutant and *35S::miR319a x 35S::miR3TCP* stable transgenic *A. thaliana* (Col-0) lines and *35S::SAP11_MBSP_* transgenic lines phenocopy only the *A. thaliana brc1 brc2* mutant. Nine-week-old plants were phenotyped for rosette leaf morphology (D), overall appearance of side views (E), rosette diameters (F) and numbers of primary branches emerging from the rosettes (G). (F, G) Error bars denote standard errors (n=24). Letters indicate groups that are statistically different (one-way ANOVA with Tukeýs Multiple Comparison Test).

To investigate which region of TCP domain determine SAP11 binding specificity, chimeras of the basic region and helix loop helix regions of the TCP domains of CIN-TCP AtTCP2 and CYC/TB1-TCP BRC1 (AtTCP18) were constructed (Fig. 2) and tested for interactions with the two SAP11 proteins in yeast two-hybrid analyses. SAP11_AYWB_ and SAP11_MBSP_ interacted with the TCP domains of AtTCP2 and BRC1 (Fig. 2B), as observed for full length TCPs (Fig. 1B), confirming that the TCP domain itself is sufficient for SAP11 interaction and specificity. Furthermore, SAP11_AYWB_ interacted with all AtTCP2-BRC1 chimeras used in the assay (Fig. 2), whereas SAP11_MBSP_ interacted with chimeras containing BRC1 helix-loop-helix and AtTCP2 basic regions, but not with those composed of AtTCP2 helix-loop-helix and BRC1 basic region or with mixed helix, loop and helix sequences (Fig. 2). Therefore, the entire helix-loop-helix region of the TCP domain is required for the specific binding of SAP11_MBSP_ to CYC/TB1 TCPs.

**Fig. 2.**
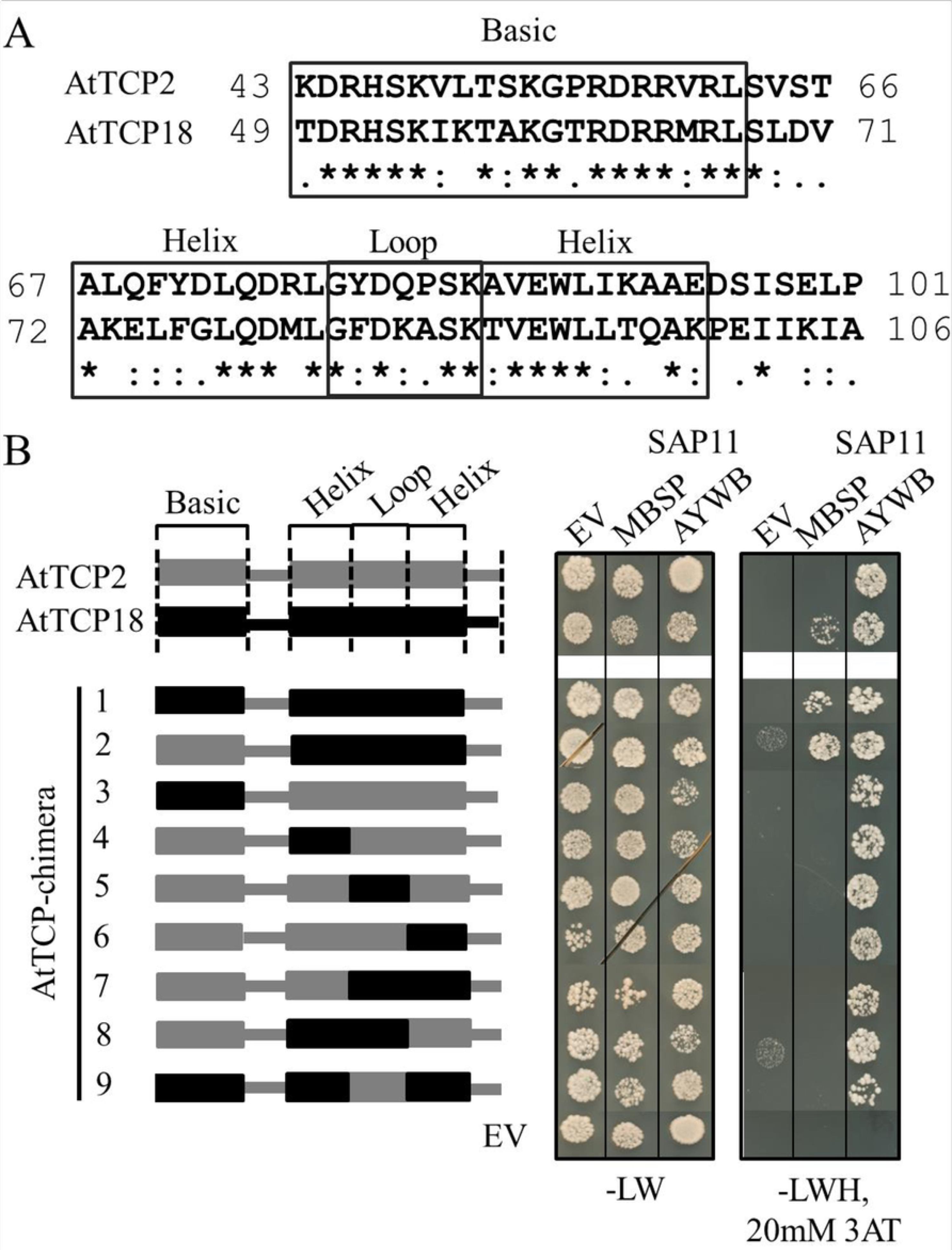
SAP11 binding specificity to regions within the TCP domains of *A. thaliana* TCP2 (CIN-TCP) and TCP18 (CYC/TB1-TCP BRC1). (A) Aligned amino acid sequences of the TCP domains of TCP2 and TCP18 with boxed basic and helix-loop-helix domains. (B) Binding specificity of SAP11_MBSP_ requires the complete TCP18 helix-loop-helix domain. Schematic representation at left are the 59-amino-acid TCP2 and TCP18 TCP domain chimeras and AtTCP2 and AtTCP18 wildtype TCP domains tested for SAP11_AYWB_ or SAP11_MBSP_ binding in yeast two-hybrid analysis at right, as described in the Fig. 1 legend.

### *A. thaliana* plants stably expressing *SAP11_MBSP_* and *SAP11_AYWB_* phenocopy *brc1 brc2* mutant or CIN-TCP knock down lines

To investigate if the SAP11 binding specificity to TCPs aligns with *in planta* interactions, phenotypes of *A. thaliana* Col-0 stable transgenic lines that produce SAP11_AYWB_ and SAP11_MBSP_ under control of the 35S promoter (Fig. 1C) were compared to those of the *A. thaliana brc1-2 brc2-1* double mutant, hereafter referred to as the *brc1 brc2* mutant, which is a *null* mutant for both CYC/TB1-TCPs BRC1 and BRC2 [34] and the *35S::miR319a x 35S::miR3TCP* line in which CIN-TCPs are knocked down [30]. Whereas the crinkled leaves of *35S::SAP11_AYWB_* lines phenocopied those of *35S::miR319a x 35S::miR3TCP* (Fig. 1D) [8], leaves of *35S::SAP11_MBSP_* lines were not crinkled and more similar to WT Col-0 leaves (Fig. 1D). Rosette diameters of the *35S::SAP11_AYWB_* and *35S::miR319a x 35S::miR3TCP* lines were smaller than WT Col-0 plants, unlike the rosettes of *35S::SAP11_MBSP_* and *A. thaliana brc1 brc2* mutant lines that looked similar to those of WT plants (Fig. 1F). Both *35S::SAP11_AYWB_* and *35S::SAP11_MBSP_* lines produced significantly more primary rosette-leaf branches (RI) [34] than WT plants. With exception of the *35S::SAP11_MBSP_* line 3 that had a lower number of RIs, the production of RI was similar to the *A. thaliana brc1 brc2* mutant. In contrast, *35S::miR319a x 35S::miR3TCP* plants produced a reduced number of RI compared to WT Col-0 (Figs. 1E and 1G, S1E Fig.). Therefore, *35S::SAP11_MBSP_* lines phenocopied the *A. thaliana brc1 brc2* mutant and the *35S::SAP11_AYWB_* lines both the *A. thaliana brc1 brc2* and *35S::miR319a x 35S::miR3TCP* mutant lines, indicating that SAP11_AYWB_ destabilizes Arabidopsis CIN and CYC/TB1 TCPs and SAP11_MBSP_ only the CYC/TB1-TCPs BRC1 and BRC2, in agreement with the results of protoplast-based destabilization and Y2H binding assays.

Beyond phenotypes described above, we found that the *35S::miR319a x 35S::miR3TCP* and *35S::SAP11_AYWB_* lines produced less rosette leaves compared to WT plants, unlike the *A. thaliana brc1 brc2* and *35S::SAP11_MBSP_* lines (S1A Fig.). Bolting time, plant height and numbers of primary cauline-leaf branches (CI) [34] were variable among the *35S::SAP11_AYWB_* and *35S::SAP11_MBSP_* lines (S1B-S1E Figs.). Roots of *35S::miR319a x 35S::miR3TCP* and *35S::SAP11_AYWB_* lines were consistently shorter compared to WT plants as described by Lu *et al*. [41]. In contrast, the root length of *A. thaliana brc1 brc2* and *35S::SAP11_MBSP_* lines did not show obvious differences compared to those of WT plants (S2 Fig.).

### SAP11_AYWB_ impairs *A. thaliana* defence responses to *M. quadrilineatus* in contrast to SAP11_MBSP_

We previously showed that the AY-WB insect vector *M. quadrilineatus* produces 20-30% more progeny on *35S::SAP11_AYWB_ A. thaliana* [8]. By repeating this experiment and including *35S::SAP11_MBSP_ A. thaliana*, we confirmed the previous result for *35S::SAP11_AYWB_ A. thaliana* but not for *35S::SAP11_MBSP_ A. thaliana* (Fig. 3A). Therefore, SAP11_AYWB_ appears to modulate plant defences in response to *M. quadrilineatus*, whereas SAP11_MBSP_ does not. To test this further, the transcriptomes of wild type, *35S::SAP11_AYWB_* and *35S::SAP11_MBSP_ A. thaliana* with and without exposure to *M. quadrilineatus* were compared via RNA-seq (S1 Table, GEO accession GSE118427). PCA showed that, in samples exposed to *M. quadrilineatus*, *35S::SAP11_MBSP_* and WT Col-0 group together, whereas the *35S::SAP11_AYWB_* samples form a separate group (Fig. 3B). Therefore, SAP11_AYWB_ has a measurable impact on the transcriptome of *A. thaliana*, unlike SAP11_MBSP_.

**Fig. 3.**
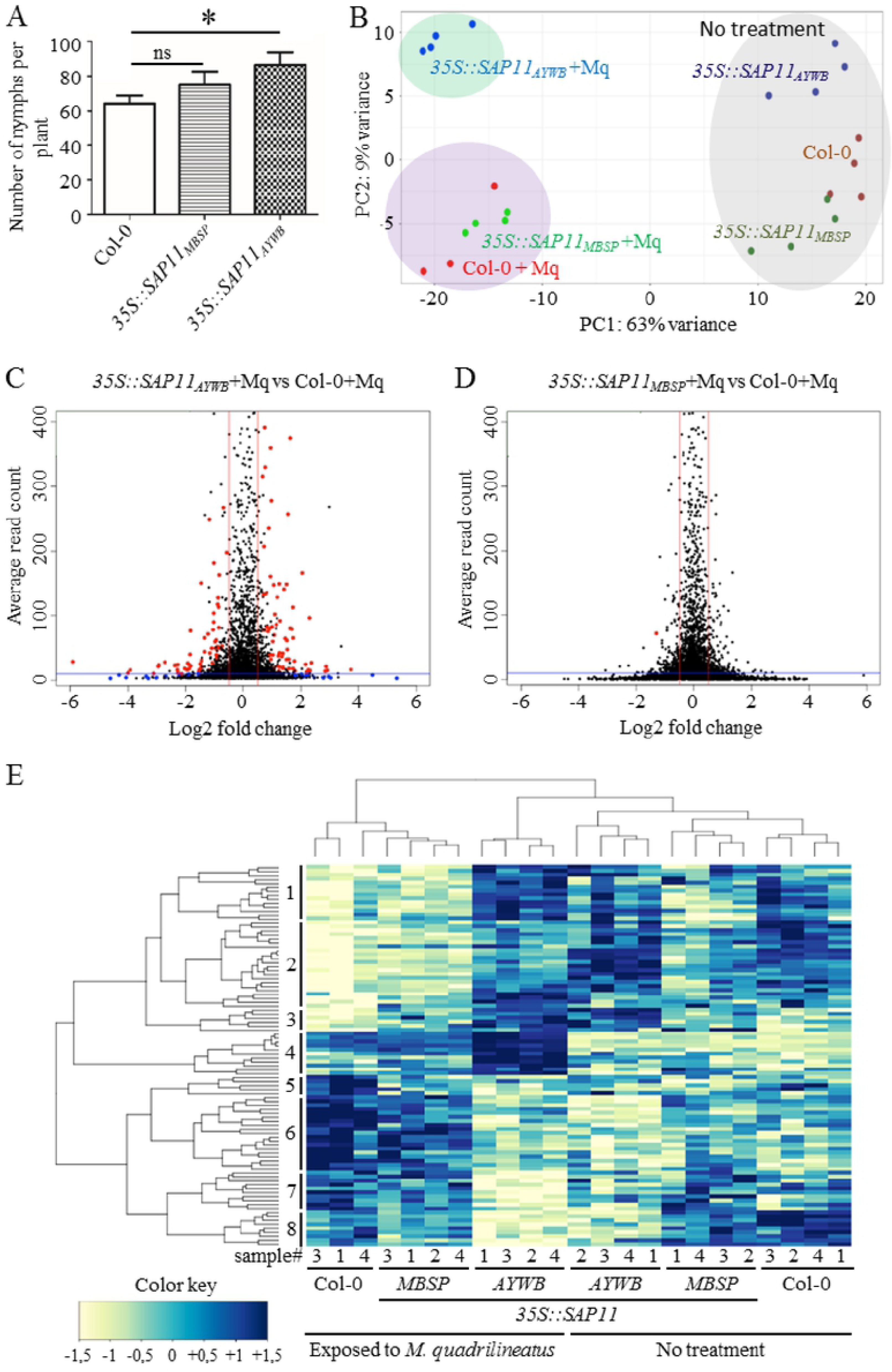
Analyses of the impact of phytoplasma SAP11_AYWB_ and SAP11_MBSP_ effectors on *A. thaliana* susceptibility to the AY-WB insect vector *M. quadrilineatus*. (A) SAP11_AYWB_ promotes *M. quadrilineatus* nymph production on *A. thaliana*, whereas SAP11_MBSP_ does not. Error bars denote standard errors, *p<0.01, students t-test compared to Col-0, n=3. (B) Principal component analysis (PCA) on the matrix of normalized read counts of 6 treatments (n=3-4, see S1 Table) showing that SAP11_AYWB_ modulates plant responses to *M. quadrilineatus* (+Mq) differently compared to SAP11_MBSP_ and wt *A. thaliana* (Col-0). (C, D) Volcano plots showing differentially expressed genes (DEGs) in insect exposed Sap11_AYWP_ and SAP11_MBSP._ DEGs with potential relevance in SAP11 dependent response (red dots) to *M. quadrilineatus* were selected by three criteria (i) P value > 0.05 (red and blue dots), (ii) average read count > 10 (dashed horizontal line) and (iii) log2 fold change > 1 (dashed vertical lines). (E) SAP11_AYWB_ modulates plant defence responses to *M. quadrilineatus* relatively to Col-0, unlike SAP11_MBSP_. Hierarchical clustering based on normalized read counts of 96 selected DEGs (red dots in C). See S2 Table for normalized read count values of all treatments and S3 Table for gene annotations with 30 genes known to be involved in defence highlighted in yellow. All experiments were executed with *35S::SAP11_AYWB_* line 7 [8] and *35S::SAP11_MBSP_* line 1 (this work).

Analyses of differentially expressed genes (DEGs) of Col-0 and transgenic plants exposed to *M. quadrilineatus* identified 96 DEGs for *35S::SAP11_AYWB_* versus Col-0 and only one DEG for *35S::SAP11_MBSP_* versus Col-0 (Figs. 3C and 3D). Hierarchical cluster of the DEGs expression levels was in agreement with the PCA results demonstrating that the *M. quadrilineatus*-exposed 35S::SAP11_AYWB_ treatments cluster separately from those of Col-0 and 35S::SAP11_MBSP_ (Fig. 3E, S2 Table). Moreover, *M. quadrilineatus*-exposed 35S::SAP11_AYWB_ treatments cluster together with non-exposed samples. Of the 96 DEGs 30 have a role in regulating plant defence responses, including hormone and secondary metabolism, such as Myb, AP2/EREBP and bZIP transcription factors, receptor kinases, cytochrome P450 enzymes, proteases, oxidases and transferases (highlighted in yellow, S3 Table). The 96 genes also included 11 natural anti-sense genes and at least 30 genes with unknown functions. Taken together, these data indicate that defence responses to *M. quadrilineatus* are suppressed in 35S::SAP11_AYWB_ plants.

### Identification of maize TCP transcription factors

To investigate SAP11 interactions with maize TCPs we first identified maize TCP sequences. The CDS of 44 *Z. mays* (Zm) TCPs available on maize TFome collection [42] were extracted from the Grass Regulatory Information Server (GRASSIUS) (http://grassius.org/grasstfdb.html) [43]. We identified two class II CYC/TB1-TCPs, including TB1 and ZmTCP18, 10 class II CIN-TCPs and 17 class I PCF-like TCPs. The ZmTCPs were assigned to groups based on characteristic TCP domain amino acids conserved in each of the groups, highlighted in yellow, red and green (Fig. 4) [28]. In contrast to *A. thaliana*, maize appears to have an additional group of class II TCPs that share amino acids conserved in the TCP domains of both CIN and TB1/CYC TCPs (Fig. 4). One of these is BRANCHED ANGLE DEFECTIVE1 (BAD1), which is expressed in the pulvinus to regulate branch angle emergence of inflorescences, particularly the tassel [44]. BAD1 was placed in a subclade of CYC-TB1 TCPs named as TCP CII. Hence, we assigned all members in this additional group to TCP CII. TCPs similar to TCP CII appear to be absent in the monocots rice (*O. sativa*) and sorghum (*S. bicolor*) (S3 and S4 Figs., S4 Table). Seven CIN-TCPs of maize, rice and sorghum are potentially regulated by miR319a (Fig. 4, S3-S5 Figs). While this study was ongoing, Chai *et al.* [45] reported the expression characteristics of 29 maize TCPs. To promote consistency, we adopted their nomenclature for these TCPs as ZmTCP01 to ZmTCP29, and continued the numbering of the additional 15 maize TCP genes extracted from GRASSIUS as ZmTCP30 to ZmTCP45 (Fig. 4, S4 Table).

**Fig. 4.**
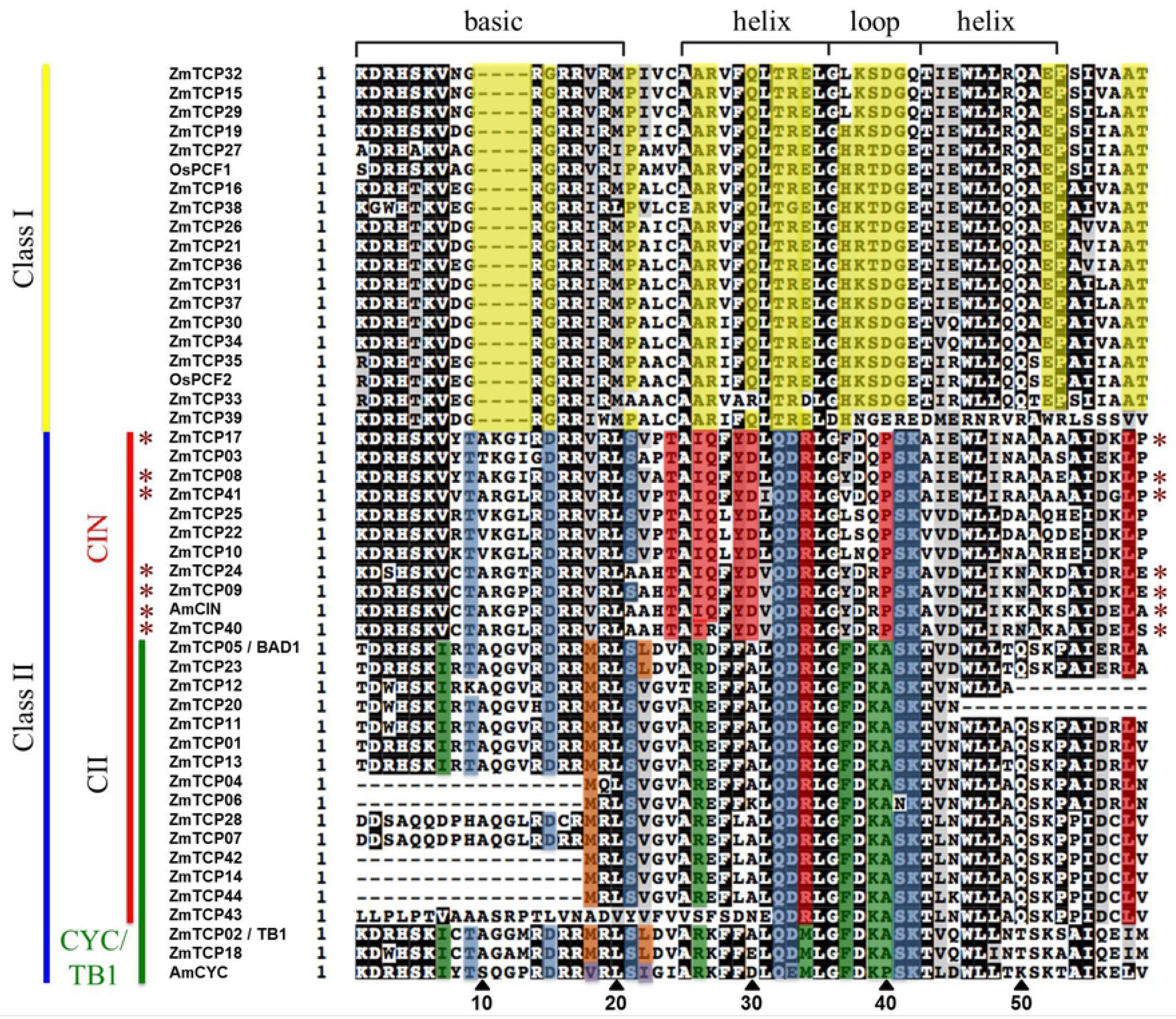
Classification of *Z. mays* (Zm) TCPs. The TCP motifs identified in 44 ZmTCPs (http://grassius.org/grasstfdb.html) were aligned with subgroup specific TCPs from *Oryza sativa* (Os) OsPCF1/2, *Antirrhinummajus* CINCINNATA (AmCIN) and CYCLOIDEA (AmCYC) and *Z. mays* TEOSINTE BRANCHED1 (ZmTCP02/TB1) (CYC/TB1 green). A number of proteins carry truncated TCP motifs at their N- or C-terminus (ZmTCP04, ZmTCP06, ZmTCP12, ZmTCP14, ZmTCP20, ZmTCP42 and ZmTCP44) or incomplete versions of the TCP-motif within their amino acid sequence (ZmTCP07, ZmTCP28, ZmTCP43). The ZmTCPs were assigned to the (sub)groups based on amino acid conservations (Class I, yellow; Class II, blue; CIN, red and CYC/TB1, green with AmCYC-like TCPs in purple and TB1-like TCPs in orange) [28] A new CII subgroup shares sequence homology with CIN-TCPs and CYC/TB1-TCPs. Asterisks indicate TCPs with potential miR319a target sites identified in their coding gene sequences (S5 Fig.).

### Phytoplasma SAP11 homologs interact with and destabilize maize class II TCPs

Y2H assays revealed that SAP11_MBSP_ interacts with the CYC/TB1-TCPs ZmTCP02 (TB1) and ZmTCP18, but not with ZmTCP members of the CIN and CII subgroups (Fig. 5A). In contrast, SAP11_AYWB_ interacted also with CIN and CII ZmTCPs (Fig. 5A). GFP-SAP11_MBSP_ and GFP-SAP11_AYWB_ destabilized HA-tagged ZmTCP02 (TB1) and ZmTCP18 in maize protoplasts in contrast to GFP controls (Fig. 5B), indicating that the SAP11 homologs also destabilize maize TCPs in maize cells.

**Fig. 5.**
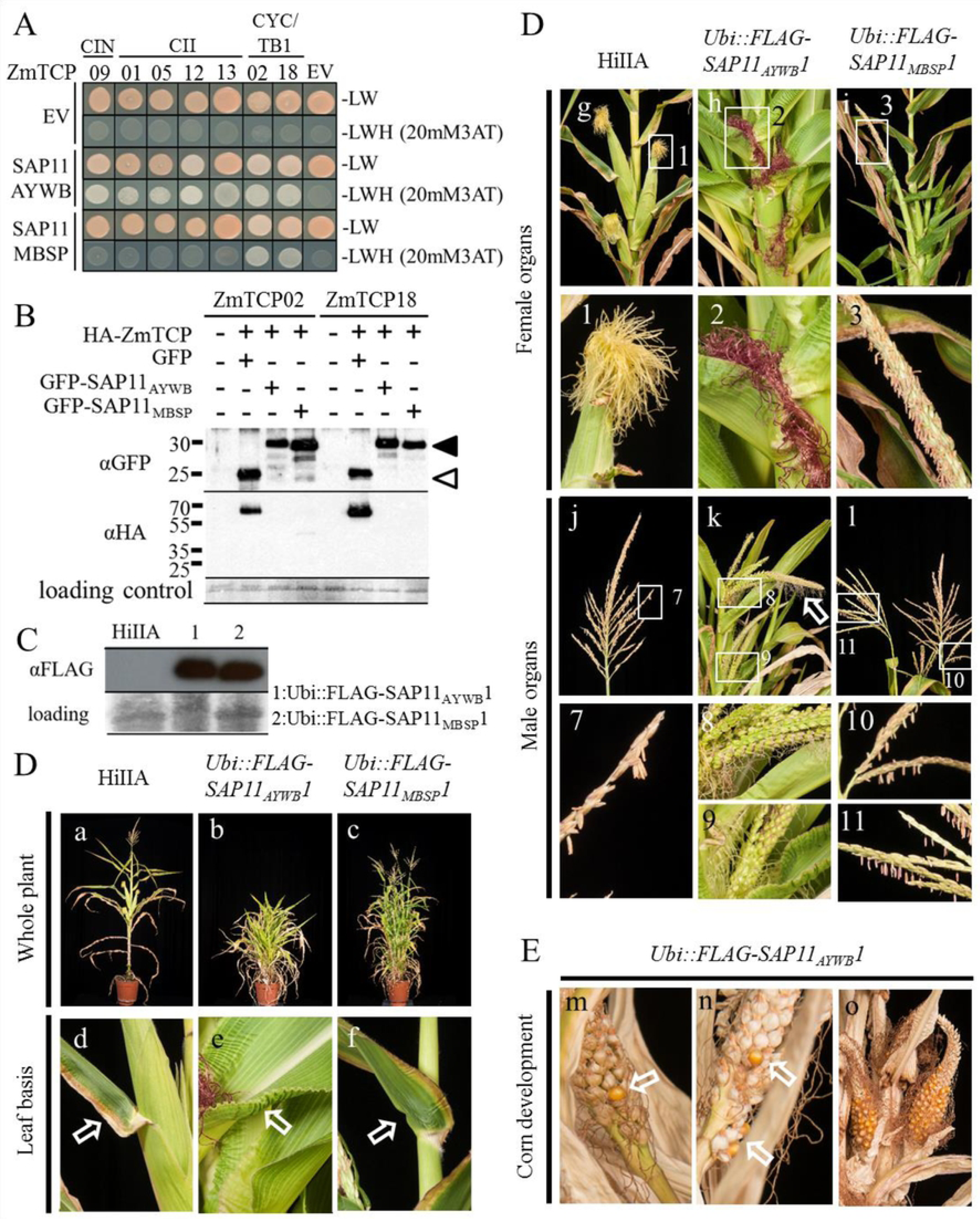
SAP11_AYWB_ and SAP11_MBSP_ interactions with maize TCP transcription factors (ZmTCPs). (A) SAP11_AYWB_ interacts with ZmTCPs of the three Class II subgroups and SAP11_MBSP_ with CYC/TB1 ZmTCPs in yeast two-hybrid (Y2H) experiments. Y2H experiments were executed with full-length ZmTCP proteins, except ZmTCP09 for which the DNA sequence corresponding to the 59 amino-acid of the TCP-motif was synthesized (Genscript). (B) SAP11_AYWB_ and SAP11_MBSP_ destabilize ZmTCPs inside maize protoplasts. Immunoblots show detection of GFP-tagged SAP11 (filled arrowheads) or GFP alone (open arrowheads) and HA-tagged TCPs with specific antibodies to GFP and HA, respectively, as indicated at left of the blots. Loading control: amidoblack staining of the large RUBISCO subunit. (C) For phenotyping FLAG-SAP11_AYWB_ (lane 1) and FLAG-SAP11_MBSP_ (lane 2) were detected in plants of the heterozygous transgenic Ubi::FLAG-SAP11_AYWB_1 and Ubi::FLAG-SAP11_MBSP_1 maize lines. The immunoblots shown were probed with anti-flag antibodies. (D) Severe developmental phenotypes of *Ubi::FLAG-SAP11_AYWB_* (HiIIA) and *Ubi::FLAG-SAP11_MBSP_* (HiIIA) transgenic maize plants. Phenotyping was done on 13-week-old transgenic and WT HiIIA plants; for each transgenic Ubi::FLAG-SAP11 maize line 3 plants were analysed and photos of one representative plant are shown. (a-c) Both *SAP11* transgenic lines are shorter and produce more tillers surrounding the main culm compared to WT HiIIA and *SAP11_MBSP_* lines also produced more axillary branches. (d-f) Crinkling of leaf edges at the base of only the *SAP11_AYWB_* lines. (g-i and insets 1-3) Impaired female inflorescence development of both *SAP11* transgenic lines. Red silk like structures emerged from the leaf sheath in the *SAP11_AYWB_* line (h, inset 2) whereas long axillary branches tipped by tassels emerged in the *SAP11_MBSP_* line (i, inset 3), compared to ears in WT HiIIA (g, inset 1). *SAP11_MBSP_* plants produced fertile pollen from these tassels, but were female sterile. (j-l, insets 7-11) Impaired male inflorescence development of *SAP11* transgenic lines. *SAP11_AYWB_* lines developed feminized tassel, including the development of silks, at the tip of the main culm (k, inset 8) and at the tip of the tillers (k, inset 9). The tassel development of *SAP11_MBSP_* lines at the tip of the main culm (l, inset 10) and at the tip of tillers (l, inset 11) resembled those of WT HiIIA (j, inset 7). (E) Feminized tassels of *SAP11_AYWB_* lines are fertile. Pollination of feminized tassels (k, insets 8 and 9) with pollen from *SAP11_MBSP_* or WT plants produced kernels (m and n), which germinated (not shown). In addition, pollination of the silks emerging from the leaf sheath (h, inset 2) resulted in the development of naked ears, without husk leaves, emerging directly from the leaf sheath (o). The ears produced kernels (o) that germinated (not shown). The *SAP11_MBSP_* lines did not produce pollen and therefore are male sterile.

### Stable *SAP11_MBSP_* and *SAP11_AYWB_* transgenic maize plants lack female and male sex organs, respectively

*SAP11_AYWB_* and *SAP11_MBSP_* were cloned as N-terminal 3XFLAG tag fusions downstream of the maize Ubiquitin promoter, and transformed into HiIIAXHiIIB hybrid *Z. mays*. *Ubi::FLAG-SAP11_MBSP_* primary transformants (T_0_) were female sterile, but produced pollen, which were used for fertilizing flowers of a wild type HiIIA plant. In contrast, *Ubi::FLAG-SAP11_AYWB_* primary transformants were male sterile, but produced flowers, which were successfully fertilized with pollen from a HiIIA plant. The T_1_ progenies of both crosses had similar production of SAP11 proteins (Fig. 5C) and were further phenotyped.

Unlike WT HiIIA, *Ubi::FLAG-SAP11_MBSP_* T_1_ plants produced multiple tillers arising from the base of the main culm (Figs. 5D (a, c) and 6). Both main culm and tillers produced apical male inflorescences with tassels that carried anthers with pollen (Figs. 5D (j, l, insets 7, 10, 11) and 6). These pollen were fertile, as they were used to pollinate HiIIA female inflorescence for seed reproduction. At the upper nodes of the main culm where in WT plants short primary lateral branches with apical ears would develop from the leaf sheath (Figs 5D (g) and 6), long primary lateral branches emerged that also had apical tassels (Figs 5D (i, inset 3) and 6). Hence, *Ubi::FLAG-SAP11_MBSP_* plants were female sterile. These phenotypes of *Ubi::FLAG-SAP11_MBSP_* plants are similar to those of the *Z. mays tb1* mutant (Fig. 6) [39,46]. Essentially, *Ubi::FLAG-SAP11_MBSP_* and *Z. mays tb1* mutant lines resemble teosinte, though the latter produces small ears located at multiple lateral positions of the primary lateral branches (Fig. 6) [47]. Therefore, *Ubi::FLAG-SAP11_MBSP_* plants phenocopy the maize *tb1* mutant, in agreement with results of yeast two-hybrid and protoplast destabilization assays showing that SAP11_MBSP_ destabilizes CYC/TB1 TCPs.

**Fig. 6.**
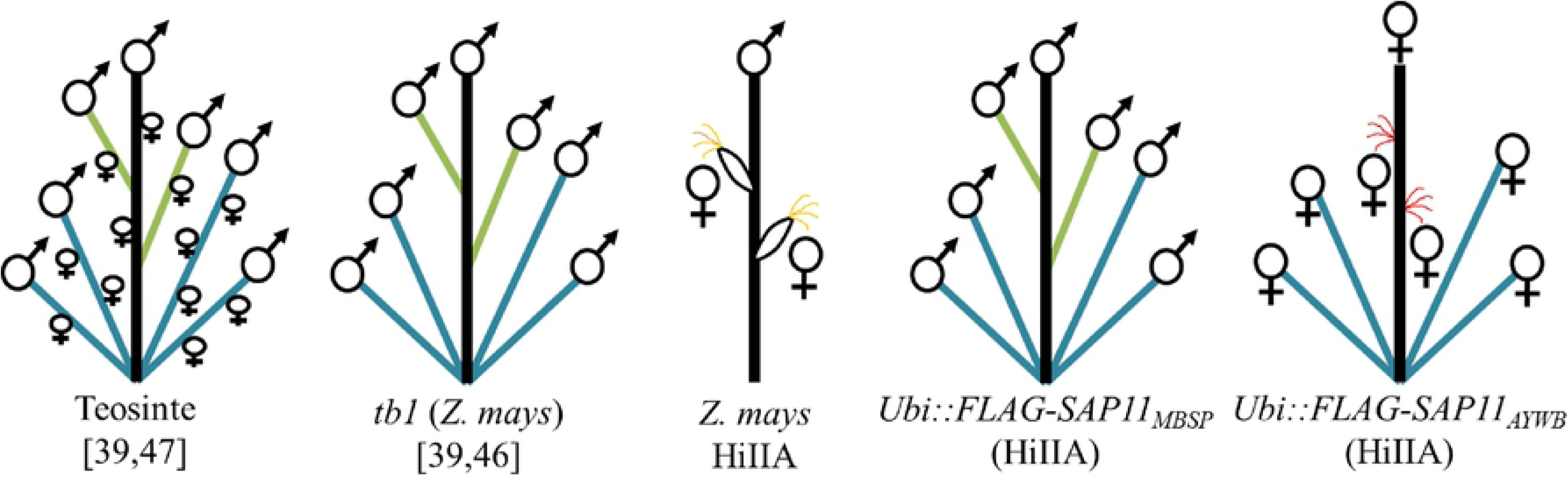
Schematic presentation of phenotyping results of WT maize (*Z. mays*), *tb1* maize and *Ubi::FLAG-SAP11* maize plants. Schematic presentations of the phenotypes of teosinte and *tb1* are included as comparison [39,46,47]. *tb1* resembles teosinte architecture but has impaired development of female inflorescences. *Ubi::FLAG-SAP11_MBSP_* plants phenocopy *tb1* plants. *Ubi::FLAG-SAP11_AYWB_* plants produce more tillers with female infloresences and naked ears from the main culm, and are male sterile. Main culms are indicated in black, axillary branches in green, tillers in blue, silks directly emerging from the main culm in red, silks of ears in yellow and inflorescences in symbols (♂, male; ♀, female).

*Ubi::FLAG-SAP11_AYWB_* T_1_ plants also produced more tillers from the base of the main culm, but were shorter than WT HiIIA and *Ubi::FLAG-SAP11_MBSP_* (Fig. 5D (a, b, c)). The majority of leaves of *Ubi::FLAG-SAP11_AYWB_* plants had curly edges, unlike *Ubi::FLAG-SAP11_MBSP_* and HiIIA plants (Fig. 5D (d, e, f, h, inset 2)). *Ubi::FLAG-SAP11_AYWB_* plants produced red-coloured silks emerging directly from the leaf sheath without prior ear formation (Figs. 5D (h, inset 2) and 6). Upon pollination of the red-coloured silks, ears with reduced husk leaves and exposed corn emerged (Fig. 5E (o)). As well, the tip of the main culm and tillers carried tassel-like structures with female flowers and emerging silks (Figs. 5D (k, insets 8, 9) and 6). Pollination of these silks with HIIA pollen induced the formation of a few corns (Fig. 5E (m,n)). Thus, *SAP11_AYWB_* induces tassel feminization and interferes with leaf development, including the modified leaves that generate the husk of the ear.

### SAP11_AYWB_ or SAP11_MBSP_ do not alter maize susceptibility to *M. quadrilineatus* and *D. maidis*

We investigated if SAP11_AYWB_ and SAP11_MBSP_ modulate maize processes in response to the AY-WB and MBSP insect vectors *M. quadrilineatus* and *D. maidis*, respectively. We did not observe any differences in fecundity of both insect vectors on HiIIA, *Ubi::FLAG-SAP11_AYWB_* and *Ubi::FLAG-SAP11_MBSP_* plants (Fig. 7A and B). PCA of RNA-seq data from WT and transgenic maize plants indicate that SAP11_AYWB_ and SAP11_MBSP_ modulate maize transcriptomes with SAP11_AYWB_ having a larger effect than SAP11_MBSP_ (Fig. 7C and D, S5 and S6 Tables, GEO: GSE118427), in agreement with morphological data of the maize lines (Figs. 5 and 6). However, *M. quadrilineatus*-exposed HiIIA *Ubi::FLAG-SAP11_AYWB_* and *Ubi::FLAG-SAP11_MBSP_* maize clustered together and separately from non-exposed maize in PCA (Fig. 7C). *D. maidis* exposed maize samples grouped with the non-exposed ones (Fig. 7D), suggesting that the SAP11 homologs do not have obvious effects on transcriptome responses of maize to the insects. Moreover, *M. quadrilineatus* has a larger impact and *D. maidis* a minor impact on maize gene expression (Fig. 7C and D). Together, these data indicate that SAP11_AYWB_ and SAP11_MBSP_ do not alter maize susceptibility to *M. quadrilineatus* and *D. maidis*.

**Fig. 7.**
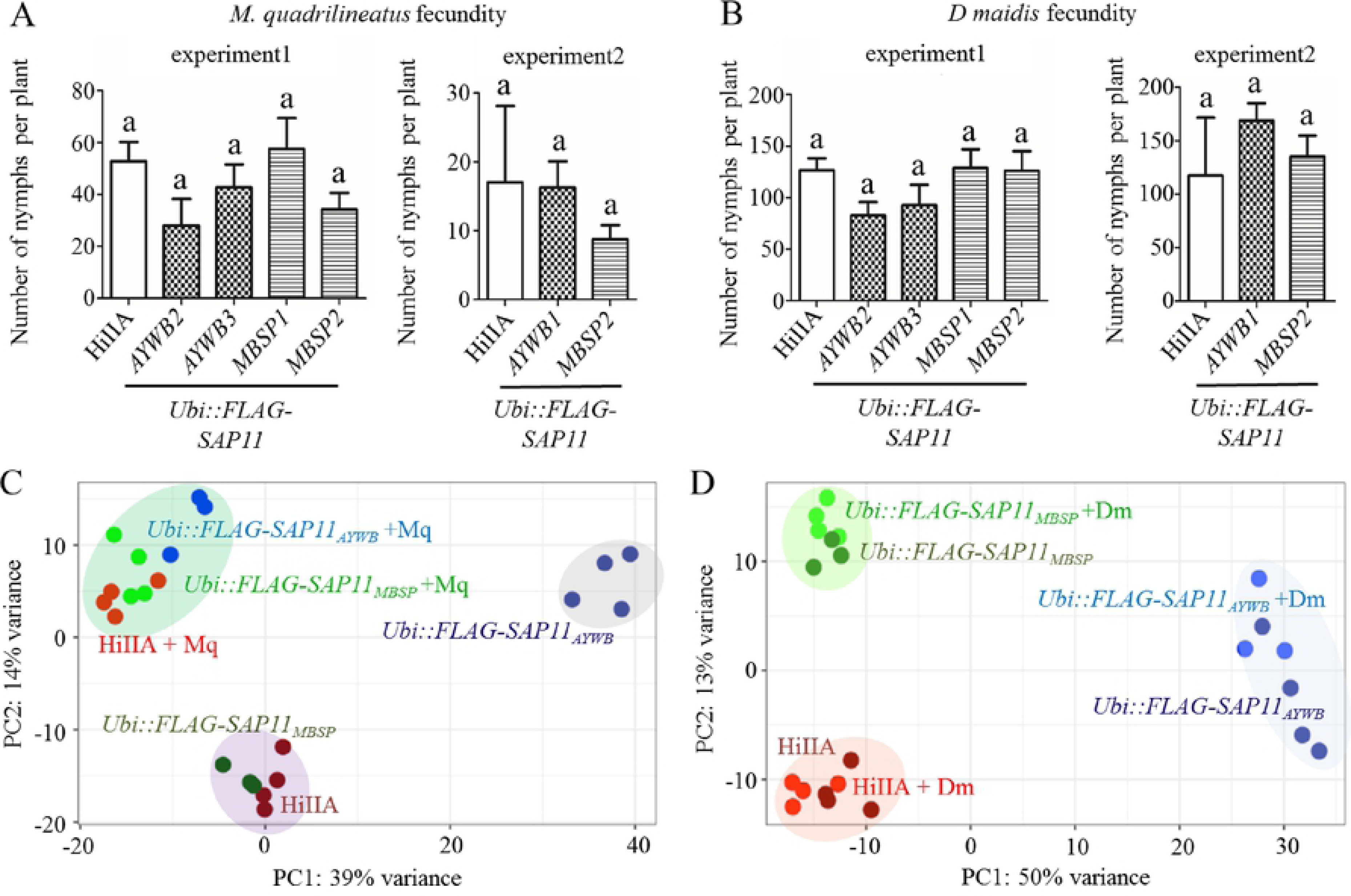
The phytoplasma SAP11_AYWB_ and SAP11_MBSP_ effectors do not modulate maize defences in response to exposure to AYWB and MBSP leafhopper vectors *M. quadrilineatus* and *D. maidis*, respectively. (A, B) Numbers of nymphs produced from the two leafhopper species are similar among *SAP11* transgenic and WT maize lines. AYWB1, 2 and 3 and MBSP1 and 2 indicate independent transgenic lines. a above the error bars indicates no significant differences (one-way ANOVA with Tukeýs Multiple Comparison Test, n=4). (C) *M. quadrilineatus* exposure (+Mq) similarly alters gene expression of *SAP11_AYWB_* and *SAP11_MBSP_* transgenic and WT maize lines. (D) Gene expression patterns of *D. maidis-*exposed (+Dm) transgenic and WT maize lines are similar to those of non-exposed lines. (C, D) Principal component analysis (PCA) on the matrix of normalized read counts of 6 treatments (n=3-4 per treatment, see S5 and S6 Tables). RNA-seq experiments were done with *Ubi::FLAG-SAP11_AYWB_* line 1 and *Ubi::FLAG-SAP11_MBSP_* line 1.

## Discussion

We found that SAP11_AYWB_ and SAP11_MBSP_ have overlapping, but distinct, binding specificities for class II TCP transcription factors. The two effectors bind to the TCP domain helix-loop-helix region. This region is required for TCP-TCP dimerization and configuration of the TCP domain beta sheets of both TCP transcription factors in a way that allows binding of the beta sheets to promoters [27]. We also found that SAP11-TCP binding specificities are correlated with the ability of the SAP11 homologs to destabilize these TCPs in leaves [8] and protoplasts (this study) and the induction of specific phenotypes in plants [8, this study]. Whereas it remains to be resolved how SAP11 destabilizes TCPs, it is clear that SAP11 is highly effective at destabilizing TCPs in plants as evidenced by the specific SAP11-induced changes in *A. thaliana* and maize architectures that phenocopy TCP mutants and knock-down lines of these plants.

TCP domains of each TCP (sub)class have characteristic amino acid sequences that have remained conserved after the divergence of monocots and eudicots [48]. We found that SAP11 binding specificity is determined by TCP (sub)class rather than plant species, as SAP11_MBSP_ specifically interacts with class II CYC/TB1-TCPs of both *A. thaliana* and maize, and not class II CIN-TCP and class I TCPs of these divergent plant species. Similarly, SAP11_AYWB_ interacts with all class II TCPs and not the class I TCPs of *A. thaliana* and maize. Therefore, SAP11_AYWB_ and SAP11_MBSP_ binding specificity is likely to involve amino acids within the helix-loop-helix region of the TCP domain that are characteristic for each TCP (sub)class and are conserved among plants species, including dicots and monocots.

We found that SAP11_MBSP_ specifically interacts with and destabilizes TCPs of the TB1 clade, including *A. thaliana* BRC1 and BRC2 and maize TCP02 and TCP18. These binding specificities are supported by plant phenotypes; *A. thaliana 35S::SAP11_MBSP_* and maize *Ubi::FLAG-SAP11_MBSP_* lines phenocopy *A. thaliana brc1 brc2* lines and maize *tb1* lines, respectively. The *A. thaliana 35S::SAP11_MBSP_* lines show stem proliferations, in agreement with *A. thaliana* BRC1 and BRC2 and maize TB1 (ZmTCP02) being suppressors of axillary bud growth [37, 49–51]. We also show that *A. thaliana 35S::SAP11_MBSP_* and *brc1 brc2* lines produce fully fertile flowers, whereas maize *Ubi::FLAG-SAP11_MBSP_* plants produced only male tassels and no female inflorescences like maize tb1 plants [39,46]. This is in agreement with BRC1 not directly affecting *A. thaliana* flower architecture [52,53], and maize TB1 being a direct positive regulator of MADS-box transcription factors that control maize female inflorescence architecture [40]. Interestingly, many phytoplasmas have SAP54 effectors, which degrade MADS-box transcription factors leading to the formation of leaf-like sterile flowers [9,10,54,55] whereas no effector gene with sequence similarity to SAP54 was identified in MBSP [56]. It is possible that the maize-specialist phytoplasma strain does not require an additional effector (such as SAP54) to modulate floral development of its host, as SAP11_MBSP_ indirectly targets flowering via TB1.

Whereas SAP11_MBSP_ interacts and destabilizes TB1 TCPs, SAP11_AYWB_ interacts with all class II TCPs of *A. thaliana* and maize, in agreement with *A. thaliana 35S::SAP11_AYWB_* lines phenocopying both *A. thaliana brc1 brc2* and *A. thaliana 35S::miR319a x 35S::miR3TCP* lines. Information about the role of TCPs in maize development are limited, potentially due to redundant functions of TCPs belonging to the same subgroup and the challenges of obtaining multiple knockdown lines. Therefore, at this time we do not know if *maize Ubi::FLAG-SAP11_AYWB_* lines phenocopy maize mutant lines for all CIN and CII TCPs. Nonetheless the leaf crinkling phenotypes of *Ubi::FLAG-SAP11_AYWB_* maize plants are in agreement with what is known about the functions of CIN TCPs in Arabidopsis where CIN TCPs play a role in leaf development [8,32,57]. The CII subgroup member BAD1 regulates branch angle emergence of the maize tassel [44] indicating that CII TCPs regulate male inflorescence development in maize. Our finding that *Ubi::FLAG-SAP11_AYWB_* maize plants solely producing female inflorescences and no tassels expands the current knowledge about maize CII and CIN-TCPs to a potential role in plant sex determination. We cannot fully exclude the possibility that SAP11_AYWB_ destabilizes other proteins in maize, though we think this is unlikely given our finding that SAP11-TCP interactions are specific involving conserved TCP helix-loop-helix sequences and that SAP11_AYWB_ induces changes in *A. thaliana* development that are entirely consistent with destabilization of class II TCPs in this plant. Therefore, phenotypes seen of *Ubi::FLAG-SAP11_AYWB_* maize plants are likely caused by SAP11_AYWB_-mediated destabilization of all maize class II TCPs, indicating a direct role of these TCPs in the development of maize male and female inflorescence architectures.

We previously demonstrated that 35S::SAP11_AYWB_ *A. thaliana* plants are affected in jasmonate production and *LOX2* expression upon wounding and that the AY-WB insect vectors produce more progeny on *LOX2*-silenced plants [8]. A number of TCPs have roles in plant JA production regulation [31, 58–63]. Here, we show a clear role of SAP11_AYWB_ suppression of plant defence response genes to *M. quadrilineatus*, including those involved in phytohormone responses. These genes were not differentially regulated in SAP11_MBSP_ plants response to *M. quadrilineatus*, indicating that destabilization of CIN-TCPs alone or in combination with Arabidopsis BRC1 and BRC2 alters plant defence responses to *quadrilineatus*. SAP11_AYWB_ does not promote *M. quadrilineatus* and *D. maidis* fecundity on maize suggesting that maize class II TCPs do not play a major role in regulating defence responses of maize leaves. Therefore, class II TCPs appear to regulate plant defence responses in leaves of Arabidopsis but not in maize.

MBSP and the insect vectors *D. maids* and *D. elimatus* are thought to have co-evolved with maize since its domestication from teosinte [23]. We previously sequenced the genomes of MBSP isolates from geographically distant locations and found single nucleotide polymorphisms (SNPs) throughout the genomes of these isolates but that SAP11_MBSP_ remained conserved [56]. The effector may be subject to purifying selection because the destabilization of maize TB1 TCPs and subsequent induction of axillary branching and inhibition of female flower production promote MBSP fitness in maize in a manner that is so far unknown. As well, SAP11_MBSP_ evolution may be constrained by possibly negative effects of maize CIN and ECE TCP destabilization on MBSP fitness or because SAP11_MBSP_ alleles that destabilize other maize TCPs may not be selected in MBSP populations because maize TCPs do not impact *D. maidis* fitness. Finally, both *D. maidis* and MBSP predominantly colonize maize, whereas *M. quadrilineatus* and AYWB colonize a wide range of plants species presenting the possibility that a positive effect of SAP11 on insect fecundity may have more benefit for a generalist phytoplasma and insect vector than for more specialized ones.

In conclusion, we found that SAP11 effectors of AY-WB and MBS phytoplasmas have evolved to target overlapping but distinct class II TCPs of their plant hosts and that these transcription factors also have overlapping but distinct roles in regulating development in these plant species. In addition, TCPs may or may not impact plant defence responses to phytoplasma leafhopper vectors. The distinct roles of TCPs in regulating plant developmental and defence networks are likely to shape SAP11 effector evolution of phytoplasma.

## Material and Methods

### Generation of Gateway^TM^ compatible entry clones

We generated Gateway^TM^ compatible entry clones for all experiments, except for the constructs to transform maize. The cloning of the codon-optimized version of SAP11_AYWB_ without the sequence corresponding to the signal peptide into pDONR207 is described previously [8]. The cloning of sequences corresponding to the open reading frames (ORFs) of AtTCP2, AtTCP3, AtTCP4, AtTCP5, AtTCP7, AtTCP10, AtTCP13 and AtTCP17 (S4 Table) into pDONR207 was also done previously [7]. The full-length ORF of AtTCP6, AtTCP8, AtTCP9, AtTCP12, AtTCP14 and AtTCP18 (S4 Table) were PCR amplified from complementary DNA (cDNA) with gene-specific primers that contain partial sequences of the attB1 and attB2 Gateway^TM^ recombination sites (S7 Table). The fragments were further amplified with attB1 and attB2 adapter primers and cloned into pDONR207 with Gateway^TM^ BP Clonase II Enzyme Mix (Invitrogen, Carlsbad, USA). Gateway^TM^ compatible pENTR/SD/D/TOPO vectors containing the full length ORFs of ZmTCP01 (clone UT5707), ZmTCP02 (clone UT5978), ZmTCP05 (clone UT1680), ZmTCP12 (clone UT6182), ZmTCP13 (clone UT3439) and ZmTCP18 (clone UT4097) were ordered from The Arabidopsis Information Resource (TAIR) (S4 Table). A codon-optimized version of SAP11_MBSP_ without the sequence corresponding to the signal peptide and DNA sequences corresponding to the TCP domains of ZmTCP9, AtTCP12, AtTCP18 and the AtTCP chimeras were gene synthesized by Genscript (New Jersey, USA) with Gateway^TM^ compatible attL1 and attL2 attachment sites (S4 and S8 Tables) and provided in pMS (Genscript).

### Transient expression assays in *Arabidopsis thaliana* and maize (Zea mays L.) protoplasts

All genes were transferred from the Gateway^TM^ compatible entry clones into the respective expression vectors with the Gateway^TM^ LR Clonase II enzyme mix (Invitrogen). Full-length ORFs of all TCPs were cloned into pUGW15 [64] to produce N-terminally HA-tagged proteins. The codon-optimized versions of SAP11_AYWB_ and SAP11_MBSP_ without signal peptide sequences were cloned into pUBN-GFP-DEST [65] to produce N-terminally GFP-tagged SAP11_AYWB_ and SAP11_MBSP_. To generate a plasmid for expression of GFP alone, the ccdB cassette of pUBN-GFP-DEST was replaced with a GFP sequence that carries two translational stop codons instead of the translational start codon. The GFP-sequence was amplified from pUBN-GFP-DEST with the gene-specific primers STOP-GFP forward and reverse (S7 Table), cloned into pDONR207 with the Gateway^TM^ BP Clonase II Enzyme Mix (Invitrogen) and transferred to pUBN-GFP-DEST using the Gateway^TM^ LR Clonase II Enzyme Mix (Invitrogen).

Isolation and transformation of Arabidopsis and maize protoplasts were performed as described by [66]. Protoplasts were generated from 6-week-old Arabidopsis and four-leaf stage maize plants grown in controlled environmental conditions with a 14h, 22 C°/ 10h, 20°C light / dark period. The maize plants were transferred into dark for five days before protoplast isolation. 600-µl-protoplast-suspensions were transformed with the indicated constructs and placed in the dark for 12h for gene expression. Protoplasts were harvested by mild centrifugation (1 min, 200 × g) and mixed with 20µl 2X sodium dodecyl sulfate (SDS)-polyacrylamide gel electrophorese (PAGE) sample buffer (50mM Tris/HCl, 10% (w:v) SDS, 50% (v:v) glycerol, 0.02% bromophenolblue, 10% ß-mercaptoethanol, pH=6.8). Samples were separated in an SDS-PAGE using 15% SDS-polyacrylamide gels and blotted on 0.45µm BA85 Whatman® Protran® nitrocellulose membranes (Sigma-Aldrich) with the BioRad (Life Science, Hemel Hempstead, UK) minigel and blotting system. Proteins were detected via western blot hybridization with specific antibodies. For detection of GFP-fusion proteins, anti-GFP polyclonal primary antibody (Santa Cruz Biotechnology, Dalla, USA, catalog number: sc-8334, diluted 1:1000) and anti-rabbit-HRP secondary antibody (Sigma-Aldrich, diluted 1:10000) were used. After the anti GFP-antibodies were removed by treatment of the membrane with 0.2 M glycine, 0.1% SDS, 100 mM ß-mercaptoethanol, pH=2, the HA-fusion proteins were detected on the same blot with anti-HA11 monoclonal primary antibody (Covance, New Jersey, USA, order number: MMS-101P, diluted 1:1000) and anti-mouse-HRP secondary antibody (Sigma-Aldrich, diluted 1:10000).

### Yeast Two-Hybrid analyses

All genes were transferred from the above generated Gateway^TM^ compatible entry clones into the respective Yeast Two-Hybrid vectors with the Gateway^TM^ LR Clonase II enzyme mix (Invitrogen). The codon-optimized sequences corresponding to mature proteins (without signal peptides) of SAP11_AYWB_ and SAP11_MBSP_ were transferred into pDEST-GAD-T7 [67]. The TCP sequences encoding for full length TCPs or TCP domains were transferred into the pDEST-GBK-T7 [67]. *Saccharomyces cerevisiae* strain AH109 (Matchmaker III; Clonetech Laboratories, Mountain View, CA, USA) was transformed using a 96-well transformation protocol [68] and interaction studies were carried out on media depleted of leucine, tryptophan and histidine with addition of 20 mM 3-Amino-1,2,4-triazole (3AT) to suppress auto activation.

### Generation and analysis of transgenic *A. thaliana* lines

The generation and analysis of the 35S::SAP11_AYWB_ Arabidopsis Col-0 lines, was described previously [8]. Idan Efroni (Weizmann Institute of Science, Rehovot, Israel) provided seeds of the 35S::miR319a x 35S::miR3TCP Arabidopsis Col-0 lines described in Efroni *et al*. [30] and Pilar Cubas (Centro Nacional de Biotecnologia, Madrid, Spain) provided seeds of the *brc1 brc2* Arabidopsis Col-0 line described in Aguilar-Martinez *et al*. [34]. For generation of the 35S::SAP11_MBSP_ Arabidopsis Col-0 lines the codon optimized version of the *SAP11_MBSP_* sequence without the sequence corresponding to the signal peptide was transferred from the Gateway^TM^ compatible entry clone (described above) into the pB7WG2 binary vector using the Gateway^TM^ LR Clonase II Enzyme Mix (Invitrogen) and Arabidopsis Col-0 plants were transformed using the floral dipping method [69].

### Quantitative Real Time-PCR experiments

SAP11 transcript levels in 35S::SAP11_AYWB_ and 35S::SAP11_MBSP_ *A. thaliana* plants were quantified in mature leaves of three independent, 5-week-old plants. Total RNAs were extracted from 100 mg snap frozen *A. thaliana* leaves with TRI-reagent (Sigma Aldrich) and cDNA synthesis was performed from 0.5 µg total RNA using the M-MLV-reverse transcriptase (Invitrogen). cDNA was subjected to qRT-PCR using SYBR® Green JumpStart™ Taq ReadyMix™ (Sigma-Aldrich) in a CFX96 Touch™ Real-Time PCR Detection System (Biorad) using gene-specific primers for the SAP11-homologs and Actin 2 (AT3G18780) (S9 Table).

### Root length measurements

*A. thaliana* seeds were sterilized in 5% sodium hypochlorite for 8 minutes and washed five times with sterile water. Seeds were germinated on ½ × MS medium with 0.8% (w/v) agar. Three days after germination, seedlings were transferred to ½ × Hoagland medium [70] with 0.25 mM KH_2_PO_4_ containing 1% (w/v) sucrose and 1% (w/v) agar [41]. Plates were placed vertical to allow root growth on the agar surface. After an additional growth period of 10 days seedlings were removed from the plates individually and their root length measured using a ruler.

### Generation and analysis of transgenic maize lines

Codon optimized versions of the *SAP11_AYWB_* and the *SAP11_MBSP_* sequences without sequences corresponding to the signal peptide including a sequence encoding an N-terminal 3xFLAG-tag were synthesized with flanking BamH1 and EcoRI restriction sites (S10 Table) that were used for cloning into the multiple cloning site of the p1u Vector (DNA Cloning Service, Hamburg, Germany). The resulting *Ubi::FLAG-SAP11-nos* cassette was transferred from p1U into the binary Vector p7i (DNA Cloning Service, Hamburg, Germany) via SfiI restriction sites. Agrobacterium-mediated transformation of maize HiIIAxHiIIB embryos was performed by Crop Genetic Systems (CGS) UG (Hamburg, Germany). T_0_ transgenic HiIIAxHiIIB plants were selected with BASTA (Bayer CropScience, Monheim, Germany). For seed reproduction T_0_ transgenic plants were crossed with HiIIA plants because the described defects in sexual organs development (Fig. 5) impeded self-pollination. Plants were analyzed for production of proteins from transgenes via western blot hybridizations (explained above) with anti-FLAG M2 monoclonal primary antibody (Sigma-Aldrich, order number: F3165, diluted 1:1000) and anti-mouse-HRP secondary antibody (Sigma-Aldrich, diluted 1:10000) and then used for experiments.

### Insect fecundity assays

Plants were grown under controlled environmental conditions with a 14h, 22 C°/ 10h, 20°C light / dark period for Arabidopsis and 16h, 26°C/ 8h, 20°C light/dark period for maize. Seven-week-old Arabidopsis and three-week-old maize plants were individually exposed to 10-15 adult *M. quadrilineatus* or *D. maidis* insects (7-10 females and 3-5 males) for 3 days. The insects were removed and progeny (nymphs or adults) were counted four weeks later.

### RNA-seq analysis

Fully expanded leaves of seven-week-old *A. thaliana* Col-0 wt and transgenic plants were exposed to five adult *M. quadrilineatus* (2 males and 3 females) in a single clip cage with one clip-cage per plant. For the generation of non-treated samples, clip-cages were applied without insects. After 48h the areas covered by the clip-cages were harvested, snap frozen in liquid nitrogen and stored at −80C until further processing for RNA extraction. For maize, complete three-week-old maize HiIIA wild type (WT) or transgenic plants were exposed to 50 adult *M. quadrilineatus* or *D. maidis* insects (20 males and 30 females) for 48 hours and the complete above soil plant material was harvested, snap frozen in liquid nitrogen and stored at −80C until further processing for RNA extraction.

Total RNA was extracted from ground Arabidopsis leaf tissue and from 200 mg ground maize material using the RNeasy plant mini kit with on-column DNase digestion (Qiagen). The RNA-seq data of the *A. thaliana* experiments were generated at Academia Sinica (Taipei, Taiwan) and at the Earlham Institute (EI, Norwich, UK). The RNA-seq data of all maize experiments were generated at EI. At Academia Sinica, libraries were generated with the llumina Truseq strand-specific mRNA library preparation without size selection, and sequenced on the Illumina HiSeq2500, 125-bp paired-end reads (YOURGENE Bioscience, New Taipei City, Taiwan). Libraries at EI were generated using NEXTflex directional RNA library (HT) preparation (Perkin Elmer, Austin, Texas, USA) and sequencing was done on the Illumina HiSeq4000, 75-bp paired-end reads (EI). To assess if the RNA-seq data for the *A. thaliana* experiments received from EI and Academia Sinica are comparable, four samples were sequenced at both facilities. Principal Component Analysis (PCA) showed that the samples generated by these two facilities cluster together demonstrating that batch effects are negligible (S6 Fig.).

The adapter sequences of the raw RNAseq reads were removed using Trim Galore, version 0.4.4 (https://www.bioinformatics.babraham.ac.uk/projects/trim_galore/). The paired-end reads were aligned to the reference genome (*A. thaliana*/TAIR 10.23 and *Z. mays*/AGPv4) with the software TopHat, version 2.1.1 [71]. The number of aligned reads per gene was calculated using HTSeq, version 0.6.1 [72], and data were initially analysed via PCA, using the R/Bioconductor package DESeq2 [73]. Obvious outliers were excluded from the analysis; this amounted to one sample per experiment, as follows: one wild type (WT) Col-0 + *M. quadrilineatus* sample from the *A. thaliana* experiment; one *Ubi::FLAG-SAP11_AYWB_* + *M. quadrilineatus* sample from one of the maize experiments; one *Ubi::FLAG-SAP11_AYWB_* + *D. maidis* sample from the other maize experiment; and one *Ubi::FLAG-SAP11_MBSP_* sample in common with both experiments (S7 Fig., S1, S5, S6 Tables). Differential expression analysis was conducted with DESeq2, using the function -contrast- to make specific comparisons. For further analyses we selected genes that satisfy 3 criteria: *p* value <0.05 after accounting for a 5% false discovery rate (FDR) (Benjamini-Hochberg corrected), mean gene expression value >10 and fold change in expression >2. Cluster analysis was performed on z-score normalized data using the hierarchical method [74].

### Transcriptome assemblies of *M. quadrilineatus* and *D. maidis* RNA-seq data

RNA-seq data of *M. quadrilineatus* and *D. maidis* males and females (∼25 million reads each) were downloaded from NCBI, accession number SRP093182 and SRP093180 respectively. The reads were used for *de novo* assemblies of male and female transcriptomes separately. Reads were trimmed to remove adaptor sequence and low-quality reads using Trim Galore (https://www.bioinformatics.babraham.ac.uk/projects/trim_galore/). Reads over 20-bp in length were retained for downstream analysis. Trimmed reads were *de novo* assembled using Trinity r20140717 [75] allowing a minimum contig length of 200 bp and minimum k-mer coverage of 2 with default parameters. Assembled contigs were made non-redundant and lowly expressed contigs were filtered with FPKM cut-off 1 using build-in Perl script provided by Trinity. This resulted in 48474 transcripts for male *M. quadrilineatus*, 44409 transcripts for female *M. quadrilineatus*, 42815 transcripts for male *D. maidis* and 59131 transcripts for female *D. maidis*. These assemblies were used to validate the origin of RNA-seq data by assessing if reads aligning to leafhopper transcripts were present in RNA-seq data derived from plants exposed to the leafhoppers as opposed to those of plants that were not exposed to the leafhoppers.

## Acknowledgments

We acknowledge Dr. Idan Efroni (Department of Plant Sciences, Weizmann Institute of Science, Rehovot, Israel) for providing seeds for the *35S::miR319a x 35S::miR3TCP* Arabidopsis Col-0 line [30], Dr. Pilar Cubas (Department of Plant Molecular Genetics, Centro Nacional de Biotecnologia, Madrid, Spain) for seeds of the *brc1 brc2* Arabidopsis Col-0 line [34], Dr. Ali Al-Subhi and Prof. Abdullah Al-Saadi (Department of Crop Sciences, Sultan Qaboos University, Muscat, Oman) for assistance with yeast two-hybrid experiments. We thank Dr. Ian Bedford, Anna Jordan, Gavin Hatt and Jake Stone from the JIC entomology team for insect rearing, and Andrew Davis for photography. We also thank Dr. Dirk Becker (Crop Genetic Systems, Hamburg, Germany) for providing maize HiIIA seeds and Dr. Dirk Becker and Dr. Uta Paszkovski (Department of Plant Sciences, University of Cambridge, Cambridge, UK) for sharing their expertise in maize cultivation. For revision of the manuscript we thank Prof. Richard Immink.

## Supporting information captions

**S1 Fig. Phenotyping of transgenic *35S::SAP11_AYWB_* and *35S::SAP11_MBSP_* Arabidopsis plants.** Three independent lines overexpressing either *SAP11_AYWB_* or *SAP11_MBSP_* were analysed in comparison to Col-0, the *brc1 brc2* mutant and *35S::miR319a x 35S::miR3TCP* with regard to (A) the number of rosette leaves when first bolting buds appeared at the centre of the leaf rosette, (B) the time point of bolting buds appearance, (C) the plant height and (D) the number of primary cauline-leaf branches (CI). The number of primary rosette-leaf branches (RI) are presented in Fig. 1G of the main text. (E) Schematic presentation of Arabidopsis branching. Error bars denote standard errors (n=24). Asterisks indicate statistically significant differences compared to Col-0. (*, p<0.05, **, p<0.01, ***, p<0.001, student’s t-test); ns, not significant.

**S2 Fig. CIN-TCP destabilization affects root lengths.** (A) Roots of representative *35S::SAP11_AYWB_* and *35S::SAP11_MBSP_* mutants compared to Col-0, the *brc1 brc2* mutant and *35S::miR319a × 35S::miR3TCP* lines. (B) Root length measurements of indicated mutants compared to Col-0. Error bars denote standard errors (n=20). Asterisks indicates statistically significant difference (*, p<0.001, student’s t-test); ns, not significant.

**S3 Fig. Classification of *Sorghum bicolor* (Sb) TCPs.** The TCP motifs of 27 SbTCPs (http://grassius.org/grasstfdb.html) were aligned and assigned to the TCP (sub)groups as described in Fig. 4. Corresponding gene codes are presented in S4 Table. SbTCP4 carries a truncated TCP-motif at its C-terminus and SbTCP10 and SbTCP23 carry incomplete versions of the TCP-motif within their amino acid sequence. Sequences were aligned using ClustalW (http://www.genome.jp/tools/clustalw/) and visualized using the Boxshade software (http://www.ch.embnet.org/software/BOX_form.html). Asterisks indicate TCPs with potential miR319a target sites identified in their coding gene sequences (S5 Fig.).

**S4 Fig. Classification of *Oryza sativa* (Oz) TCPs.** The TCP motifs of 27 OzTCPs (http://grassius.org/grasstfdb.html) were aligned and assigned to the TCP (sub)groups as described in Fig. 4. Corresponding gene codes are presented in S4 Table. Sequences were aligned using ClustalW (http://www.genome.jp/tools/clustalw/) and visualized using the Boxshade software (http://www.ch.embnet.org/software/BOX_form.html). Asterisks indicate TCPs with potential miR319a target sites identified in their coding gene sequences (S5 Fig.).

**S5 Fig. Identification of potential miR319a target sites.** The CDS of the TCPs from *Zea mays* (Zm), *Oryza sativa* (Os), *Sorghum bicolor* (Sb), and of the *Antirrhinum majus* (Am) CIN-TCP were screened for potential miR319a target sites. They are depicted together with the miR319a binding sites of *Arabidopsis thaliana* (At) CIN-TCPs [57]. Nucleotides known to be involved in miR319a binding to AtCIN-TCPs are indicated in grey [57].

**S6 Fig. Principal component analysis (PCA) showing that samples sequenced at different facilities cluster together (batch effect is negligible).** PCA was conducted with normalized read counts of RNA-seq data obtained from *M. quadrilineatus*-exposed leaves of three *A. thaliana* Col-0 plants (samples #1, 2 and 3) and *35S::miR319a x 35S::miR3TCP* sample #4 generated at the Earlham Institute, Norwich, UK (red circles) and Academia Sinica, Taipei, Taiwan (green triangles).

**S7 Fig. Cluster analysis performed on the matrix of normalized read counts of RNA-seq values from** (A) *Arabidopsis* Col-0, *35S::SAP11_AYWB_* and *35S::SAP11_MBSP_* non-exposed and exposed to *M quadrilineatus* (+Mq). (B) *Z. mays* HiIIA, *Ubi::FLAG-SAP11_AYWB_* and *Ubi::FLAG-SAP11_MBSP_* non-exposed and exposed to *M. quadrilineatus* (+Mq) and (C) non-exposed and exposed to *D. maidis* (+Dm). Experiments were done with *35S::SAP11_AYWB_* line 7 (Sugio *et al.*, 2011b), *35S::SAP11_MBSP_* line 1, *Ubi::FLAG-SAP11_AYWB_* line 1 and *Ubi::FLAG-SAP11_MBSP_* line 1.

**S1 Table: Alignment to SAP11 transgene, M. quadrilineatus transcriptome and A. thaliana genome of RNA-seq data shown in Fig. 3.** +Mq indicates samples from *M. quadrilineatus* exposed plants.

**S2 Table: List of differentially expressed genes and expression values in RNA-seq experiments of 6 treatments.** The genes are ordered according to the heat map in Fig 3E. Genes potentially involved in plant defense response are highlighted in yellow and annotations of these genes are listed in S3 Table. +Mq indicates samples from *M. quadrilineatus* exposed plants.

**S3 Table: List of differentially expressed genes with potential biological functions.** The genes are ordered according to the heat map in Fig 3E. Genes potentially involved in plant defense response are highlighted in yellow.

**S4 Table: Sequence IDs of TCPs from Zea mays (Zm), Arabidopsis thaliana (At), Sorghum bicolor (Sb) and Oryza sativa (Os).**

S5 Table: Alignment to SAP11 transgene, *M. quadrilineatus* transcriptome and *Z. mays* genome of RNA-seq data of *M. quadrilineatus*-exposed (+Mq) *Z. mays* shown in Fig. 7.

S6 Table: Alignment to SAP11 transgene, *D. maidis* transcriptome and *Z. mays* genome of RNA-seq data of *D. maidis*-exposed (+Dm) *Z. mays* shown in Fig. 7.

**S7 Table. Oligonucleotide sequences (5′ > 3′) for cloning.**

**S8 Table. Synthesized CDS (underlined) flanked by gateway compatible attL1 and attL2 sites.** Nucleotide sequences for gene syntheses of *SAP11_MBSP_* for expression in *Arabidopsis thalina* and of the TCP domains from *ZmTCP33*, *AtTCP2*, *AtTCP18* and chimeras of *AtTCP2* and *AtTCP18* TCP domains for expression in yeast.

**S9 Table. Oligonucleotide sequences (5′ > 3′) for qRT-PCR.**

**S10 Table. Nucleotide sequences for gene syntheses of *FLAG-SAP11_MBSP_* and *FLAG-SAP11_AYWB_* for expression in *Zea mays.*** Kodzak sequences are in italic, ORFs are flanked by *Bam*HI and *Eco*R1 restriction sites (grey) for subsequent cloning.

## References

1. Weisburg WG, Tully JG, Rose DL, Petzel JP, Oyaizu H, Yang D, et al. A phylogenetic analysis of the mycoplasmas: basis for their classification. Journal of Bacteriology. 1989;171(12):6455–67.

2. Gundersen DE, Lee IM, Rehner SA, Davis RE, Kingsbury DT. Phylogeny of mycoplasmalike organisms (Phytoplasmas): a basis for their classification. Journal of Bacteriology. 1994;176(17):5244–54.

3. Lee IM, Davis RE, Gundersen-Rindal DE. Phytoplasma: phytopathogenic mollicutes. Annu Rev Microbiol. 2000;54:221–55.

4. Weintraub PG, Beanland L. Insect vectors of phytoplasmas. Annu Rev Entomol. 2006;51:91–111.

5. Bertaccini A. Phytoplasmas: diversity, taxonomy, and epidemiology. Front Biosci. 2007;12:673–89.

6. Hogenhout SA, Oshima K, Ammar el D, Kakizawa S, Kingdom HN, Namba S. Phytoplasmas: bacteria that manipulate plants and insects. Mol Plant Pathol. 2008;9(4):403–23.

7. Sugio A, MacLean AM, Kingdom HN, Grieve VM, Manimekalai R, Hogenhout SA. Diverse targets of phytoplasma effectors: from plant development to defense against insects. Annu Rev Phytopathol. 2011;49:175–95.

8. Sugio A, Kingdom HN, MacLean AM, Grieve VM, Hogenhout SA. Phytoplasma protein effector SAP11 enhances insect vector reproduction by manipulating plant development and defense hormone biosynthesis. Proc Natl Acad Sci U S A. 2011;108(48):E1254–63.

9. MacLean AM, Orlovskis Z, Kowitwanich K, Zdziarska AM, Angenent GC, Immink RG, et al. Phytoplasma effector SAP54 hijacks plant reproduction by degrading MADS-box proteins and promotes insect colonization in a RAD23-dependent manner. PLoS Biol. 2014;12(4):e1001835.

10. Kitazawa Y, Iwabuchi N, Himeno M, Sasano M, Koinuma H, Nijo T, et al. Phytoplasma-conserved phyllogen proteins induce phyllody across the Plantae by degrading floral MADS domain proteins. J Exp Bot. 2017;68(11):2799–811.

11. Sugio A, MacLean AM, Hogenhout SA. The small phytoplasma virulence effector SAP11 contains distinct domains required for nuclear targeting and CIN-TCP binding and destabilization. New Phytol. 2014;202(3):838–48.

12. MacLean AM, Sugio A, Makarova OV, Findlay KC, Grieve VM, Toth R, et al. Phytoplasma effector SAP54 induces indeterminate leaf-like flower development in Arabidopsis plants. Plant Physiol. 2011;157(2):831–41.

13. Orlovskis Z, Hogenhout SA. A bacterial parasite effector mediates insect vector attraction in host plants independently of developmental changes. Front Plant Sci. 2016;7:885.

14. Janik K, Mithofer A, Raffeiner M, Stellmach H, Hause B, Schlink K. An effector of apple proliferation phytoplasma targets TCP transcription factors-a generalized virulence strategy of phytoplasma? Mol Plant Pathol. 2017;18(3):435–42.

15. Chang SH, Tan CM, Wu CT, Lin TH, Jiang SY, Liu RC, et al. Alterations of plant architecture and phase transition by the phytoplasma virulence factor SAP11. J Exp Bot. 2018;69(22):5389–401.

16. Wang N, Yang H, Yin Z, Liu W, Sun L, Wu Y. Phytoplasma effector SWP1 induces witches’ broom symptom by destabilizing the TCP transcription factor BRANCHED1. Mol Plant Pathol. 2018;19(12):2623–34.

17. Bai X, Zhang J, Ewing A, Miller SA, Radek AJ, Shevchenko DV, et al. Living with genome instability: the adaptation of phytoplasmas to diverse environments of their insect and plant hosts. J Bacteriol. 2006;188(10):3682–96.

18. Toruño TY, Music MS, Simi S, Nicolaisen M, Hogenhout SA. Phytoplasma PMU1 exists as linear chromosomal and circular extrachromosomal elements and has enhanced expression in insect vectors compared with plant hosts. Mol Microbiol. 2010;77(6):1406–15.

19. Chung WC, Chen LL, Lo WS, Lin CP, Kuo CH. Comparative analysis of the peanut witches’-broom phytoplasma genome reveals horizontal transfer of potential mobile units and effectors. PLoS One. 2013;8(4):e62770.

20. Ku C, Lo WS, Kuo CH. Horizontal transfer of potential mobile units in phytoplasmas. Mob Genet Elements. 2013;3(5):e26145.

21. Lee IM, Gundersen-Rindal DE, Davis RE, Bottner KD, Marcone C, Seemuller E. ‘*Candidatus* Phytoplasma asteris’, a novel phytoplasma taxon associated with aster yellows and related diseases. Int J Syst Evol Microbiol. 2004;54(Pt 4):1037–48.

22. Sugio A, Hogenhout SA. The genome biology of phytoplasma: modulators of plants and insects. Curr Opin Microbiol. 2012;15(3):247–54.

23. Nault LR, Delong DM. Evidence for co-evolution of leafhoppers in the genus Dalbulus (*Cicadellidae: Homoptera*) with maize and its ancestors. Annals of the Entomological Society of America. 1980;73(4):349–53.

24. Gonzalez JG, Jaramillo MG, Lopes JRS. Undetected infection by maize bushy stunt phytoplasma enhances host-plant preference to *Dalbulus maidis* (*Hemiptera: Cicadellidae*). Environmental entomology. 2018;47(2):396–402.

25. Navaud O, Dabos P, Carnus E, Tremousaygue D, Herve C. TCP transcription factors predate the emergence of land plants. J Mol Evol. 2007;65(1):23–33.

26. Cubas P, Lauter N, Doebley J, Coen E. The TCP domain: a motif found in proteins regulating plant growth and development. The Plant journal: for cell and molecular biology. 1999;18(2):215–22.

27. Aggarwal P, Das Gupta M, Joseph AP, Chatterjee N, Srinivasan N, Nath U. Identification of specific DNA binding residues in the TCP family of transcription factors in Arabidopsis. Plant Cell. 2010;22(4):1174–89.

28. Martin-Trillo M, Cubas P. TCP genes: a family snapshot ten years later. Trends Plant Sci. 2009;15(1):31–9.

29. Howarth DG, Donoghue MJ. Duplications in *Cys*-like genes from Dipsacales correlate with floral form. Int J Plant Sci. 2005;166(3):357–70.

30. Efroni I, Blum E, Goldshmidt A, Eshed Y. A protracted and dynamic maturation schedule underlies Arabidopsis leaf development. Plant Cell. 2008;20(9):2293–306.

31. Schommer C, Debernardi JM, Bresso EG, Rodriguez RE, Palatnik JF. Repression of cell proliferation by miR319-regulated TCP4. Mol Plant. 2014;7(10):1533–44.

32. Nicolas M, Cubas P. The role of TCP transcription factors in shaping flower structure, leaf morphology, and plant architecture. 2015:249–67.

33. Vadde BVL, Challa KR, Nath U. The TCP4 transcription factor regulates trichome cell differentiation by directly activating GLABROUS INFLORESCENCE STEMS in *Arabidopsis thaliana*. The Plant journal: for cell and molecular biology. 2018;93(2):259–69.

34. Aguilar-Martinez JA, Poza-Carrion C, Cubas P. Arabidopsis BRANCHED1 acts as an integrator of branching signals within axillary buds. Plant Cell. 2007;19(2):458–72.

35. Koyama T, Furutani M, Tasaka M, Ohme-Takagi M. TCP transcription factors control the morphology of shoot lateral organs via negative regulation of the expression of boundary-specific genes in Arabidopsis. Plant Cell. 2007;19(2):473–84.

36. Gonzalez-Grandio E, Poza-Carrion C, Sorzano CO, Cubas P. BRANCHED1 promotes axillary bud dormancy in response to shade in Arabidopsis. Plant Cell. 2013;25(3):834–50.

37. Yang Y, Nicolas M, Zhang J, Yu H, Guo D, Yuan R, et al. The TIE1 transcriptional repressor controls shoot branching by directly repressing BRANCHED1 in Arabidopsis. PLoS Genet. 2018;14(3):e1007296.

38. Luo D, Carpenter R, Copsey L, Vincent C, Clark J, Coen E. Control of organ asymmetry in flowers of Antirrhinum. Cell. 1999;99(4):367–76.

39. Doebley J., Stec A, Gustus C. *teosinte branched1* and the origin of maize: evidence for epistasis and the evolution of dominance. Genetics. 1995;141:333–46.

40. Studer AJ, Wang H, Doebley JF. Selection during maize domestication targeted a gene network controlling plant and inflorescence architecture. Genetics. 2017;207(2):755–65.

41. Lu YT, Li MY, Cheng KT, Tan CM, Su LW, Lin WY, et al. Transgenic plants that express the phytoplasma effector SAP11 show altered phosphate starvation and defense responses. Plant Physiol. 2014;164(3):1456–69.

42. Burdo B, Gray J, Goetting-Minesky MP, Wittler B, Hunt M, Li T, et al. The Maize TFome--development of a transcription factor open reading frame collection for functional genomics. The Plant journal: for cell and molecular biology. 2014;80(2):356–66.

43. Yilmaz A, Nishiyama MY, Jr., Fuentes BG, Souza GM, Janies D, Gray J, et al. GRASSIUS: a platform for comparative regulatory genomics across the grasses. Plant Physiol. 2009;149(1):171–80.

44. Bai F, Reinheimer R, Durantini D, Kellogg EA, Schmidt RJ. TCP transcription factor, BRANCH ANGLE DEFECTIVE 1 (BAD1), is required for normal tassel branch angle formation in maize. Proc Natl Acad Sci U S A. 2012;109(30):12225–30.

45. Chai W, Jiang P, Huang G, Jiang H, Li X. Identification and expression profiling analysis of TCP family genes involved in growth and development in maize. Physiol Mol Biol Plants. 2017;23(4):779–91.

46. Hubbard L, McSteen P, Doebley J, Hake S. Expression patterns and mutant phenotype of *teosinte branched1* correlate with growth suppression in maize and teosinte. Genetics. 2002;162(4):1927–35.

47. Brown PJ, Upadyayula N, Mahone GS, Tian F, Bradbury PJ, Myles S, et al. Distinct genetic architectures for male and female inflorescence traits of maize. PLoS Genet. 2011;7(11):e1002383.

48. Horn S, Pabón-Mora N, Theuß VS, Busch A, Zachgo S. Analysis of the CYC/TB1 class of TCP transcription factors in basal angiosperms and magnoliids. The Plant Journal. 2014;81(4):559–71.

49. Finlayson SA. Arabidopsis TEOSINTE BRANCHED1-LIKE 1 regulates axillary bud outgrowth and is homologous to monocot TEOSINTE BRANCHED1. Plant Cell Physiol. 2007;48(5):667–77.

50. Dong Z, Li W, Unger-Wallace E, Yang J, Vollbrecht E, Chuck G. Ideal crop plant architecture is mediated by *tassels replace upper ears1*, a BTB/POZ ankyrin repeat gene directly targeted by TEOSINTE BRANCHED1. Proc Natl Acad Sci U S A. 2017;114(41):E8656–e64.

51. Gonzalez-Grandio E, Pajoro A, Franco-Zorrilla JM, Tarancon C, Immink RG, Cubas P. Abscisic acid signaling is controlled by a BRANCHED1/HD-ZIP I cascade in Arabidopsis axillary buds. Proc Natl Acad Sci U S A. 2017;114(2):E245–E54.

52. Niwa M, Daimon Y, Kurotani K, Higo A, Pruneda-Paz JL, Breton G, et al. BRANCHED1 interacts with FLOWERING LOCUS T to repress the floral transition of the axillary meristems in Arabidopsis. Plant Cell. 2013;25(4):1228–42.

53. Niwa M, Endo M, Araki T. Florigen is involved in axillary bud development at multiple stages in Arabidopsis. Plant signaling & behavior. 2013;8(11):e27167.

54. Maejima K, Iwai R, Himeno M, Komatsu K, Kitazawa Y, Fujita N, et al. Recognition of floral homeotic MADS domain transcription factors by a phytoplasmal effector, phyllogen, induces phyllody. The Plant journal: for cell and molecular biology. 2014;78(4):541–54.

55. Maejima K, Kitazawa Y, Tomomitsu T, Yusa A, Neriya Y, Himeno M, et al. Degradation of class E MADS-domain transcription factors in Arabidopsis by a phytoplasmal effector, phyllogen. Plant signaling & behavior. 2015;10(8):e1042635.

56. Orlovskis Z, Canale MC, Haryono M, Lopes JRS, Kuo CH, Hogenhout SA. A few sequence polymorphisms among isolates of Maize bushy stunt phytoplasma associate with organ proliferation symptoms of infected maize plants. Ann Bot. 2017;119(5):869–84.

57. Palatnik JF, Edwards A, Wu X, Schommer C, Schwab R, Carrington JC, et al. Control of leaf morphogenesis by microRNAs. Nature. 2003;425.

58. Schommer C, Palatnik JF, Aggarwal P, Chetelat A, Cubas P, Farmer EE, et al. Control of jasmonate biosynthesis and senescence by miR319 targets. PLoS Biol. 2008;6(9):e230.

59. Sarvepalli K, Nath U. Hyper-activation of the TCP4 transcription factor in *Arabidopsis thaliana* accelerates multiple aspects of plant maturation. The Plant journal: for cell and molecular biology. 2011;67(4):595–607.

60. Danisman S, van der Wal F, Dhondt S, Waites R, de Folter S, Bimbo A, et al. Arabidopsis class I and class II TCP transcription factors regulate jasmonic acid metabolism and leaf development antagonistically. Plant Physiol. 2012;159(4):1511–23.

61. Zhang C, Ding Z, Wu K, Yang L, Li Y, Yang Z, et al. Suppression of Jasmonic Acid-Mediated Defense by Viral-Inducible MicroRNA319 Facilitates Virus Infection in Rice. Mol Plant. 2016;9(9):1302–14.

62. Yang L, Teixeira PJ, Biswas S, Finkel OM, He Y, Salas-Gonzalez I, et al. *Pseudomonas syringae* type III effector HopBB1 promotes host transcriptional repressor degradation to regulate phytohormone responses and virulence. Cell Host Microbe. 2017;21(2):156–68.

63. Jiao Y, Lee YK, Gladman N, Chopra R, Christensen SA, Regulski M, et al. MSD1 regulates pedicellate spikelet fertility in sorghum through the jasmonic acid pathway. Nat Commun. 2018;9(1):822.

64. Nakagawa T, Ishiguro S, Kimura T. Gateway vectors for plant transformation. Plant Biotech. 2009:275–84.

65. Grefen C, Donald N, Hashimoto K, Kudla J, Schumacher K, Blatt MR. A ubiquitin-10 promoter-based vector set for fluorescent protein tagging facilitates temporal stability and native protein distribution in transient and stable expression studies. The Plant journal: for cell and molecular biology. 2010;64(2):355–65.

66. Yoo SD, Cho YH, Sheen J. Arabidopsis mesophyll protoplasts: a versatile cell system for transient gene expression analysis. Nat Protoc. 2007;2(7):1565–72.

67. Rossignol P, Collier S, Bush M, Shaw P, Doonan JH. Arabidopsis POT1A interacts with TERT-V(I8), an N-terminal splicing variant of telomerase. J Cell Sci. 2007;120(Pt 20):3678–87.

68. Pecher P, Eschen-Lippold L, Herklotz S, Kuhle K, Naumann K, Bethke G, et al. The *Arabidopsis thaliana* mitogen-activated protein kinases MPK3 and MPK6 target a subclass of ‘VQ-motif’-containing proteins to regulate immune responses. New Phytol. 2014;203(2):592–606.

69. Logemann E, Birkenbihl RP, Ulker B, Somssich IE. An improved method for preparing Agrobacterium cells that simplifies the Arabidopsis transformation protocol. Plant Methods. 2006;2:16.

70. Hoagland DR, Arnon DI. The water-culture method for growing plants without soil. California Agri- cultural Experimental Station Circular 1950;347:1–39.

71. Kim D, Pertea G, Trapnell C, Pimentel H, Kelley R, Salzberg SL. TopHat2: accurate alignment of transcriptomes in the presence of insertions, deletions and gene fusions. Genome biology. 2013;14(4):R36.

72. Anders S, Pyl PT, Huber W. HTSeq--a Python framework to work with high-throughput sequencing data. Bioinformatics (Oxford, England). 2015;31(2):166–9.

73. Love MI, Huber W, Anders S. Moderated estimation of fold change and dispersion for RNA-seq data with DESeq2. Genome biology. 2014;15(12):550.

74. Murtagh F, Legendre P. Ward’s hierarchical agglomerative clustering method: which algorithms implement Ward’s criterion? Journal of Classification. 2014;31:274–95.

75. Grabherr MG, Haas BJ, Yassour M, Levin JZ, Thompson DA, Amit I, et al. Full-length transcriptome assembly from RNA-Seq data without a reference genome. Nature biotechnology. 2011;29(7):644–52.

